# Extracellular vesicles regulate yeast growth, biofilm formation, and yeast-to-hypha differentiation in *Candida albicans*

**DOI:** 10.1101/2021.01.21.427696

**Authors:** Leandro Honorato, Joana Feital Demetrio, Cameron C. Ellis, Alicia Piffer, Yan Pereira, Susana Frases, Glauber Ribeiro de Sousa Araújo, Bruno Pontes, Maria Tays Mendes, Marcos Dias Pereira, Allan J. Guimarães, Natalia Martins da Silva, Gabriele Vargas, Luna Joffe, Maurizio Del Poeta, Joshua D. Nosanchuk, Daniel Zamith, Flavia Coelho Garcia dos Reis, Marcio L. Rodrigues, Sharon de Toledo Martins, Lysangela Ronalte Alves, Igor C. Almeida, Leonardo Nimrichter

## Abstract

The ability to undergo morphological changes during adaptation to distinct environments is exploited by *Candida albicans* and has a direct impact on virulence. In this study, we investigated the influence of fungal extracellular vesicles (EVs) during yeast growth, biofilm formation, and morphogenesis in *C. albicans*. Addition of *C. albicans* EVs (*Ca* EVs) to the culture medium positively affected yeast growth. Using crystal violet staining and scanning electron microscopy (SEM), we demonstrated that *Ca* EVs inhibited biofilm formation by *C. albicans in vitro*. By time-lapse microscopy and SEM, we showed that *Ca* EV-treatment stops filamentation promoting pseudohyphae formation with multiple sites for yeast budding. The ability of *Ca* EVs to regulate dimorphism was further compared to EVs isolated from different *C. albicans* strains, *Saccharomyces cerevisiae*, and *Histoplasma capsulatum*. *Ca* EVs from distinct strains robustly inhibited yeast-to-hyphae differentiation with morphological changes occurring in less than 4 hours. A minor inhibitory effect was promoted by EVs from *S. cerevisiae* and *H. capsulatum* only after 24 hours of incubation. The inhibitory effect of *Ca* EVs was promoted by a combination of lipid compounds identified by gas chromatography-tandem mass spectrometry analysis as sesquiterpenes, diterpenes, and fatty acids. Remarkably, *Ca* EVs were also able to reverse filamentation, transforming hyphal growth to yeast forms. Transcriptomic analysis demonstrated that treatment with *Ca* EVs modified the expression of more than 300 genes. The most effectively upregulated pathways were related to DNA metabolism. The downregulated genes were mostly associated with extracellular and adhesion proteins. Finally, yeast cells treated with *Ca* EVs for 24 hours lost their agar invasive ability and were avirulent when inoculated in *Galleria mellonella* larvae. In summary, our results indicate that fungal EVs can profoundly modify *C. albicans* growth and regulate yeast-to-hypha differentiation inhibiting biofilm formation and virulence.

## Introduction

*Candida albicans* is a common colonizer of the human skin and mucosa ^1–3^. However, in circumstances where the epithelial barrier, the immune system and/or the microbiome are compromised, this species can overgrow and cause superficial and disseminated, life-threatening infections ^4–6^. The ability of *C. albicans* to adapt and colonize distinct organs and tissues is associated with the expression of virulence factors and fitness attributes ^4^. Among them, a major strategic feature that facilitates *C. albicans* to persist in different environments is the capacity to switch morphological stages, a tightly regulated process developed by fungal organisms called dimorphism ^7^.

Dimorphic transitions in *C. albicans* involve an extensive transcriptional reprogramming (reviewed by Basso et al ^5^) and an intense cell wall remodeling ^8^. The three morphological stages of *C. albicans* regularly found in biofilm and infected tissues are yeast, pseudohypha, and hypha ^9^. Regulation of morphological changes are closely linked to pathogenesis. For this reason, a number of studies dealing with morphogenesis in *C. albicans* have been explored and several signaling pathways regulating cell shape have been described ^5, 9^. Different environmental stimuli can induce dimorphism in *C. albicans*. For instance, yeast-to-hypha differentiation is stimulated by *N*-acetylglucosamine (GlcNAc), neutral pH, serum supplementation at 37°C, 5% CO_2_, low nitrogen and amino acids, among others ^8, 10–13^.

Dimorphism in *C. albicans* can be also controlled by quorum sensing (QS), a mechanism of microbial communication ^14, 15^. Fungal QS molecules (QSM) are released in culture media according to the cell density. After reaching a threshold QSM regulate genes involved with virulence, biofilm formation and morphogenesis ^16, 17^. Farnesol, farnesoic acid, and tyrosol are major morphogenesis-related QSM produced by *C. albicans* ^17–19^, although other alcohols derived from amino acids have been also characterized ^20^. Farnesol and farnesoic acid prevent yeast-to-hypha differentiation with no effects on growth rates ^17, 21^. Tyrosol stimulates germ tube and hyphal formation ^18^. These QSM are also able to regulate biofilm formation ^16, 22^. Furthermore, production of QSM such as farnesol and tyrosol is also described in *S. cerevisiae* and trans-species crosstalk might occur ^23^.

Since the first description of fungal extracellular vesicles (EVs) by Rodrigues et al. in 2007 ^37^, a number of studies have shown that fungal organisms use EVs as a mechanism of molecular exportation ^24–36^. Heterogeneous populations of EVs containing lipids, proteins, polysaccharides, pigments and nucleic acids cross the cell wall outwards, reaching the extracellular environment ^26, 27, 30, 32, 34, 37, 38^. Their implication during pathological and physiological processes is under investigation by a number of groups. *In vitro*, fungal EVs from *Cryptococcus neoformans, C. albicans, Aspergillus flavus* and *Paracoccidioides brasiliensis* are internalized by macrophages stimulating nitric oxide and cytokine production^30, 33, 39, 40^. EVs from *A. flavus* and *P. brasiliensis* are also able to promote M1 polarization ^39, 41^. In addition, dendritic cells treated with EVs from *C. albicans* up-regulated the expression of CD86 and MHCII ^30^. Fungal EVs were also used to stimulate the innate immune system of *Galleria mellonella* protecting the larvae stage against a subsequent infection with *C. albicans, C. neoformans* and *A. flavus* ^30, 39, 42^. In contrast, EVs can also be disease-enhancing, as demonstrated by EV exposure facilitating yeast traversal across the brain blood barrier in a murine model of cryptococcosis and increasing fungal burdens and lesion diameters in mice infected with *Sporothrix brasiliensis* ^36, 43^.

EVs also mediate communication between fungal cells. Bielska and colleagues demonstrated that EVs isolated from a virulent strain of *C. gattii* were efficiently taken up by macrophages previously infected with a non-virulent strain of the same species ^28^. Inside phagocytes the EVs promoted a rapid intracellular fungal growth transferring virulence to the non-virulent strain through a mechanism apparently dependent on intact vesicles. The presence of stable RNA and proteins was also required ^28^. EVs also have a critical role during biofilm matrix production and drug resistance in *C. albicans* ^44^.

In this work, we demonstrate that EVs produced by *C. albicans* cells (*Ca* EVs) impact yeast growth, biofilm formation, yeast-to-hypha differentiation gene expression, and virulence. Our results reveal previously unknown roles of fungal EVs that will likely impact the physiology and pathogenesis of *C. albicans*.

## Results

### EVs produced by yeast forms of *C. albicans* (*Ca* EVs) stimulate the growth of yeast cells *in vitro*

*C. albicans* usually grows as yeast forms in RPMI media under agitation at 37 °C ^45^. Using the *C. albicans* strain 90028, we initially investigated the direct effect of yeast form-derived *Ca* EVs during yeast growth under these conditions (Fig. 1A). Addition of *Ca* EVs to *C. albicans* cultures significantly increased growth in RPMI-MOPS media in a dose-dependent fashion (Fig. 1A**, upper panel**). To rule out that the effect was not related to nutrient supplementation, the yeast cells were also treated with *Ca* EVs for 24 h, centrifuged, washed 3 times with PBS, enumerated, and then transferred to fresh medium. Results showed growth increase in the presence of *Ca* EVs slightly but significantly higher than in the control (PBS); however, no EV dose-dependence was observed after 24-h incubation (Fig. 1A**, lower panel**). Only yeast cells were visualized during the experiment (data not shown).

**Figure 1.**
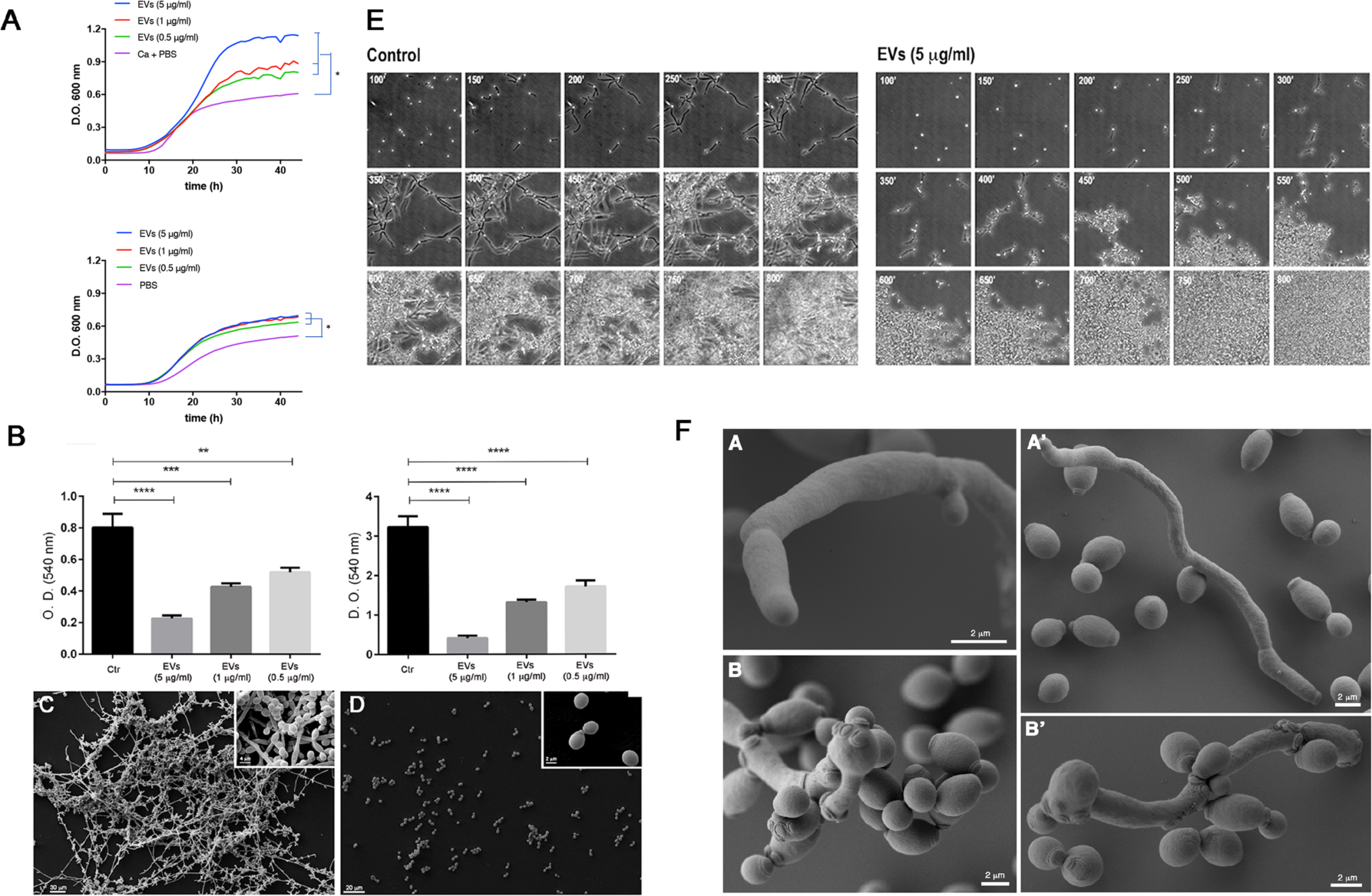
EVs from *C. albicans* stimulate yeast growth, inhibits biofilm formation and yeast-to-hyphae differentiation. Both protocols described in Methods for EV isolation were used with identical results. The results presented here were obtained from *Ca* EVs isolated from liquid culture medium. (A) Yeasts of *C. albicans* (90028) (10^3^ yeasts) were inoculated into RPMI-MOPS in the presence or absence of *Ca* EVs. Yeasts were treated with *Ca* EVs (0.5, 1, or 5 μg/ml, based on sterol content). Alternatively, treated yeasts (24 h) were washed and an aliquot was transferred to fresh medium. Cellular density was monitored for 72 h. Data shown represent one of three consistent, biological replicates. Group comparisons were submitted to one-way analysis of variance (ANOVA) with Dunnett’s correction (* p<0.0001). (B) Yeasts of *C. albicans* (90028) (10^3^ yeasts) were inoculated into RPMI-MOPS in the presence or absence of *Ca* EVs. Different concentrations of EVs (0.5, 1, and 5 μg/ml, based on sterol content), were added to wells containing *C. albicans* yeasts (strain 90028) and cells were incubated to develop biofilm for 24 and 48 h in 96-well microplates. Biofilm formation was quantified by Crystal Violet method. (C) For SEM, yeasts of *C. albicans* (strain 90028), untreated or treated with *Ca* EVs (5 μg/ml), were inoculated onto coverslips previously coated with poly-L-lysine in RPMI-MOPS and incubated for 24 h. (D) SEM images show the effect of EVs after 24 h of growth. Insets in C and D depict higher magnification of each condition. Bar, 30 μm. For both experiments, the control included culture without *Ca* EV addition. (E) Yeasts of *C. albicans* (90028) were inoculated into M199 medium to induce filamentation in the presence or absence of *Ca* EVs (5 μg/ml, based on sterol content). Cellular density was monitored, and the pictures represent the minutes of incubation (from 100 to 800 min). Growth under control conditions (absence of *Ca* EVs) and in the presence of *Ca* EVs. The numbers indicate the time in minutes. Bar, 30 μm. (F) Yeasts of *C. albicans* (90028) were inoculated into M199 medium to induce filamentation in the presence or absence of EVs (5 μg/ml, based on sterol content). SEM of cells after 24 h showing the hyphae and yeasts under control conditions (A and A’), and pseudohyphae with multiple budding sites and yeasts (B and B’).

### EVs produced by *C. albicans* inhibit biofilm formation.

Our results showed the presence of yeast-derived *Ca* EVs negatively impacted biofilm formation in a dose-dependent manner (Fig. 1B). The highest density of EVs (sterol concentration corresponding to 5 μg/ml) reduced OD measurements by approximately 80-85% when compared to controls with no *Ca* EV stimulation. SEM images obtained after 24 h of biofilm formation showed that the negative control (biofilm formed in the absence of *Ca* EVs) exhibited a standard architecture, with the presence of hypha, pseudohypha, and yeast cells attached to the coverslip (Fig. 1C). SEM images confirmed that treatment of *C. albicans* yeast cells with EVs inhibited biofilm formation. Contrasting with control conditions where the three morphological stages were found, only yeast cells were visualized attached to the coverslip in the condition with EVs present (Fig. 1D). Thus, under the conditions used in our experiments, EVs significantly impaired biofilm formation in *C. albicans*.

### EVs from *C. albicans* inhibit yeast-to-hypha differentiation

The ability to accelerate yeast growth and inhibit biofilm formation suggested that *Ca* EVs could control morphogenesis in *C. albicans* strain 90028. We then monitored microscopically the impact of *Ca* EVs on yeast-to-hypha differentiation induced by M199 medium at 37 °C. Under control conditions (absence of *Ca* EVs), the yeast cells adhered to the solid substrate (Fig. 1E**, left panel**) and formed germ tubes after 1-2 h of incubation. This step was followed by true hypha formation, which elongated with time. Few hyphae branching and yeast cells were visualized after 8 h, but hypha represented the predominant morphological stage. The time-lapse microscopy of 18 h incubation is presented as a supplemental movie (**Mov. S1**). Addition of EVs to the well drastically changed the phenotype (Fig. 1E**, right panel, and Movie S2**). Although the yeast cells were able to attach to the plate and started to form germ tubes, true hypha development after elongation was not observed (Fig. 1E**, right panel, and Movie S2**). In contrast, the appearance of constrictions on elongated cells suggested the development of pseudohyphae after 4-7 h. In addition, the cells lost their ability to adhere to the well after 6 h of incubation (**Movie S2**). Multiple budding points, potentially related to a faster growth, were observed along the pseudohyphae and after 24 h only yeast cells were detected in the presence of EVs (Figs. 1E, **right panel, and** Fig. 1F). Long pseudohyphal filaments with yeast cells clusters were observed, including a large number of yeast cells derived from pseudohyphae (Fig 1E**, right panel**). Furthermore, there was no indication of cytotoxicity under these conditions since all the yeast cells were able to grow (**Movie S2**). SEM images show the ultrastructural details of pseudohyphae after EV addition. Multiple budding yeasts and several bud scars were observed, when compared to hyphae in control condition (no *Ca* EV addition) (Fig 1F). This phenomenon of yeast induction occurred in a dose-dependent manner. A minimum density of *Ca* EVs, equivalent to 0.3-0.6 μg sterol/ml, was required to inhibit morphogenesis at least partially (Fig. 2A). However, the inhibitory activity was not dependent on the number of yeast cells, since different cell suspensions (1.5×10^3^ to 3.5×10^4^ yeast cells/well) were similarly affected using the same density (5 μg sterol/ml) of EVs (data not shown).

**Figure 2.**
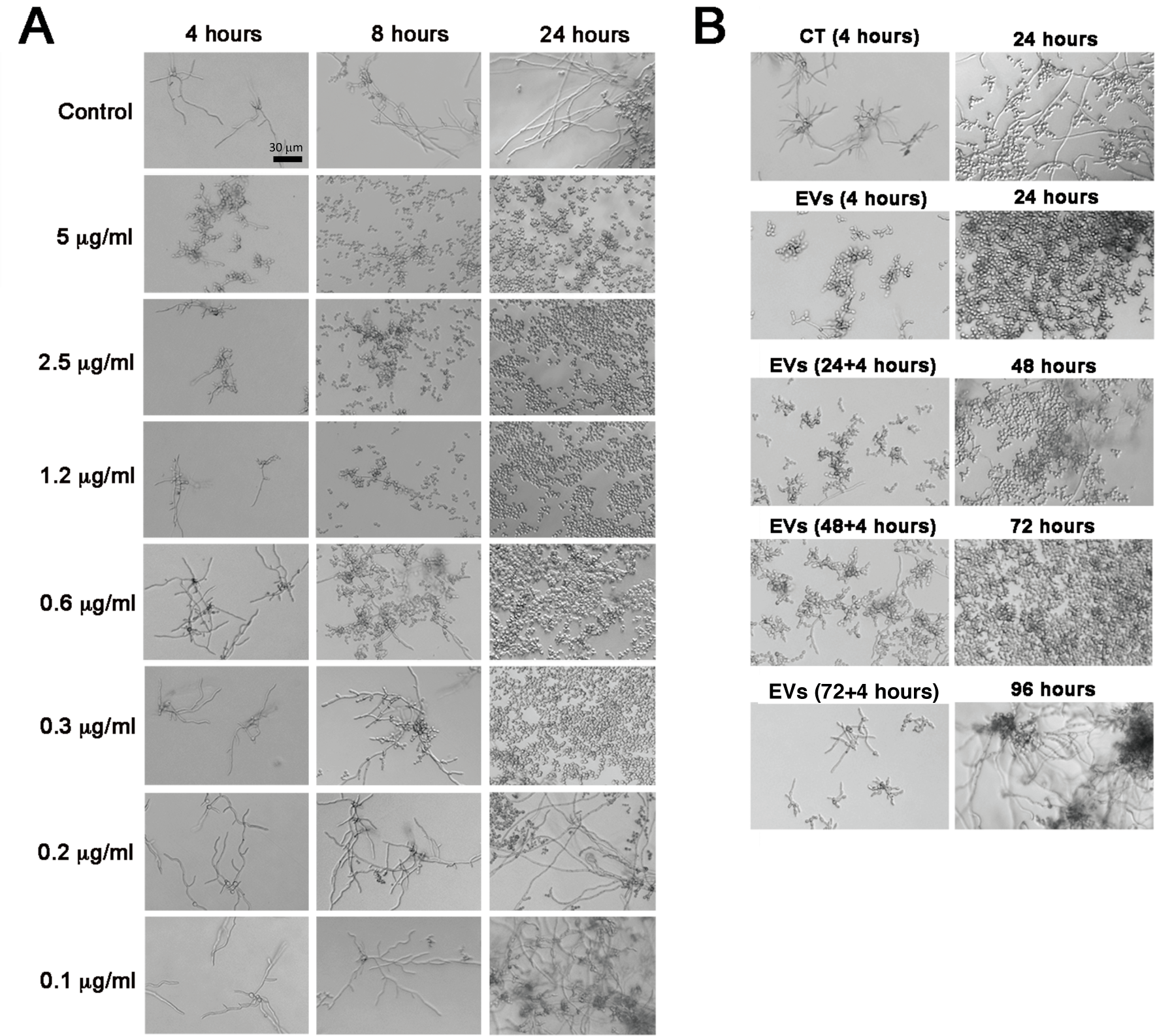
The inhibitory effect of EVs from *C. albicans* (strain 90028) on yeast-to-hypha conditions is dose-dependent and has a long-term effect. (A) Yeasts of *C. albicans* (90028) were inoculated into M199 (pH 7) in the presence or absence of *Ca* EVs in different concentrations (equivalent to 0.1, 0.2, 0.3, 0.6, 1, 1.2, 2.5, and 5 μg sterol/ml). Bar, 30 μm. (B) The inhibitory effect of a single *Ca* EV treatment was tested after 24, 48, 72, and 96 h. Aliquots of pre-treated yeasts were transferred to fresh M199 medium each 24 hours of incubation and the differentiation accompanied microscopically. Bar, 30 μm. Experiments were performed in biological triplicate with consistent results.

Remarkably, a single treatment with EVs was sufficient to inhibit the yeast-to-hypha differentiation for at least 72 h (Fig. 2B). Aliquots of pre-treated yeast cells (2.5×10^3^ yeast cells/well) were washed in PBS and plated onto fresh M199 medium and accompanied microscopically for 24 h and then transferred again to fresh medium. This process was repeated three times and aliquots from 48 and 72 h were added to fresh M199 medium for microscopic monitoring. Regular hypha formation similar to control conditions was observed only after the 96 h.

### General properties of EVs produced by different *C. albicans* strains

EVs produced by different species of *C. albicans* were isolated from culture supernatants and compared using nanoparticle tracking analysis (NTA), as described ^46^. The size distribution of the EVs produced by *C. albicans* strains used in this work ranged from 100 to 500 nm (Fig. 3A, B and C), similar to measurements published for EVs released by *C. albicans* and other fungal species ^30, 46^. EVs released by strains SC5342 and 10231 were comparable to each other, ranging from 100 to 400 nm (Fig. 3A and C). EVs produced by strain 90028 were also mainly composed by smaller particles, ranging between 100 to 250 nm, but also depicted minor populations between 300-350 nm and 420-480 nm (Fig. 3B). Considering that the *C. albicans* strains release EVs with distinct average size, we normalized our experiments using the total content of sterol in each vesicle preparation, as used in previous studies from our group and other laboratories ^30, 33^. The method is highly reproducible and requires small amounts of samples.

**Figure 3.**
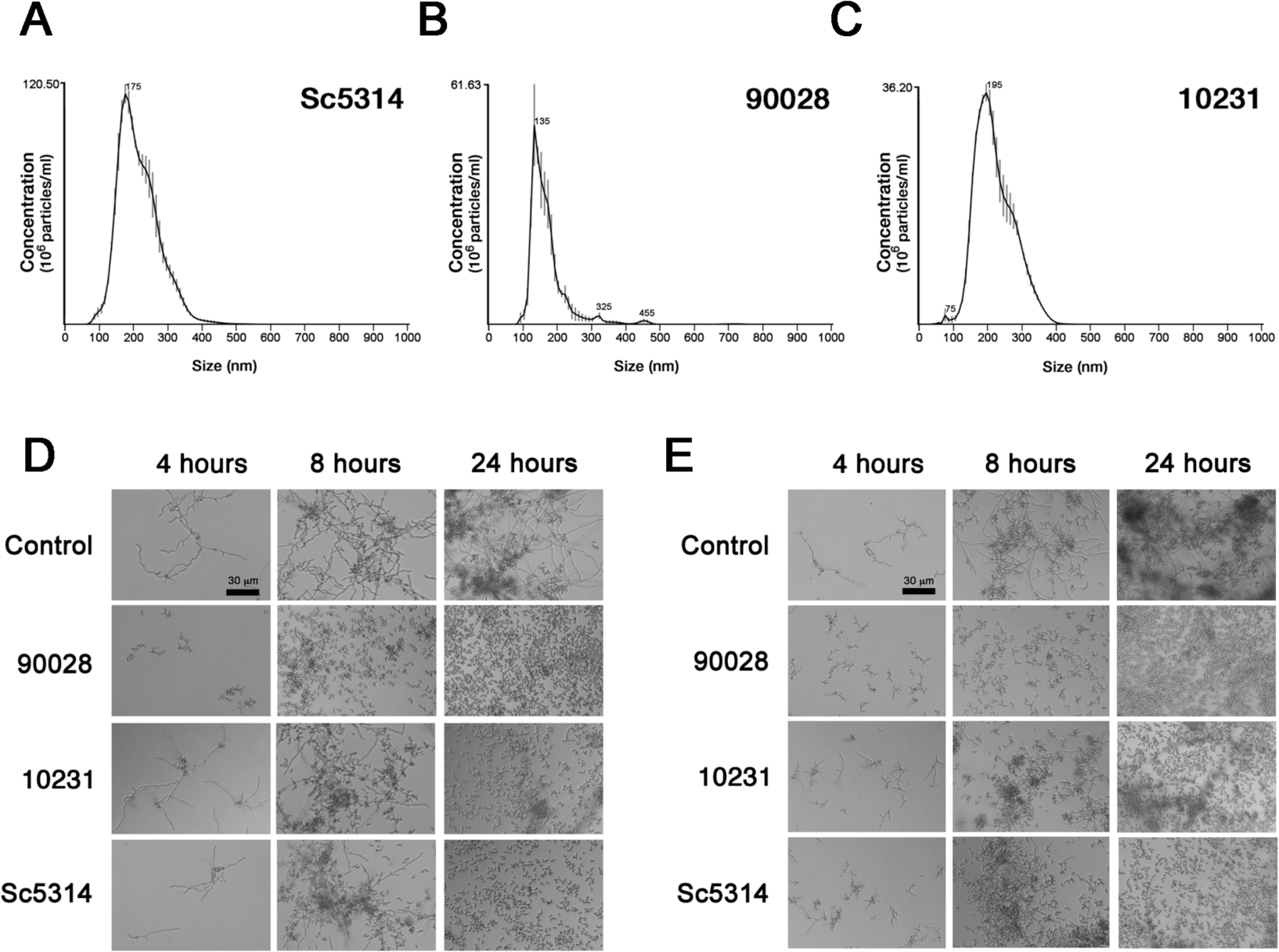
EVs from different *C. albicans* strains inhibit yeast-to-hyphae transition of strains 90028 and 10231. NTA profile of EVs produced by *C. albicans* strains (A) SC5314, (B) 90028, and (C) 10231 showed similar properties of EVs. Yeasts of *C. albicans* strains (D) 90028 and (E) 10231 were inoculated into M199 (pH 7) in the presence or absence of *Ca* EVs (equivalent to 5 μg sterol/ml) from distinct strains of *C. albicans* (90028, 10231 and SC5314). Effect was visualized after 4, 8 and 24 h. PBS was added alone as a control. Bar, 30 μm. Experiments were performed four times with consistent results.

### The ability of EVs to inhibit yeast-to-hypha differentiation is reproducible in other strains of *C. albicans*

We investigated whether the ability to inhibit yeast-to-hypha differentiation is reproducible in EVs obtained from other strains of *C. albicans*. Thus, EVs were isolated from strains with distinct abilities to cause infection. For instance, strain 10231 is less virulent and does not produce farnesol ^47^. Instead, it produces farnesoic acid, a farnesol derivative at least tenfold less active in its ability to inhibit yeast-to-hypha conversion ^21^. Strain SC5314, on the other hand, is a regular producer of farnesol ^48^. The results presented in Fig. 3 show that *Ca* EVs from both strains were able to inhibit hypha formation of strain 90028. To confirm that the effect was not strain-specific, we evaluated the inhibitory effect of the EVs produced by the *C. albicans* strains over strain 10231 (Fig. 3E). The EV-mediated inhibition of filamentation was confirmed. Further experiments were performed using EVs produced by the ATCC90028 strain in liquid medium. Although the recovery of EVs are similar using solid or liquid medium (average of 0.16 and 0.18 μg of sterol/10^9^ cells, respectively), resulting therefore in better yields of EV.

### EV molecules with yeast-to-hypha inhibitory activity are thermo-resistant and enriched in preparations containing non-polar lipids

The ability to prevent yeast-to-hypha differentiation was preserved by *Ca* EVs after heating them at 90 °C for 15 min (Fig. 4A), suggesting that thermo-resistant molecules could be mediating the inhibitory effect. Since lipids seem to be involved with *C. albicans* morphogenesis ^16, 19^, we extracted and partially fractioned the lipid components carried by *Ca* EVs and tested their inhibitory activity. The crude lipid was partitioned using standardized protocols to enrich lipids according to their polarity ^49^, and the organic (lower and upper) phases were added to yeast suspensions in M199 medium for microscopic monitoring of morphogenesis. The lower phase (LP) activity was very similar to *Ca* EVs in its ability to inhibit differentiation (Fig. 4B). Although the upper phase (UP) was also active, its effect was observed only after 24 h of incubation. The precipitate (protein-rich fraction, PF) obtained from *Ca* EV lipid extraction, likely rich in proteins, had no apparent influence on morphogenesis (Fig. 4B). The LP of lipids extracted from the whole cells were also able to inhibit *C. albicans* differentiation (data not shown). LP obtained from liquid Sabouraud using the same extraction and partition protocol showed no activity (data not shown).

**Figure 4.**
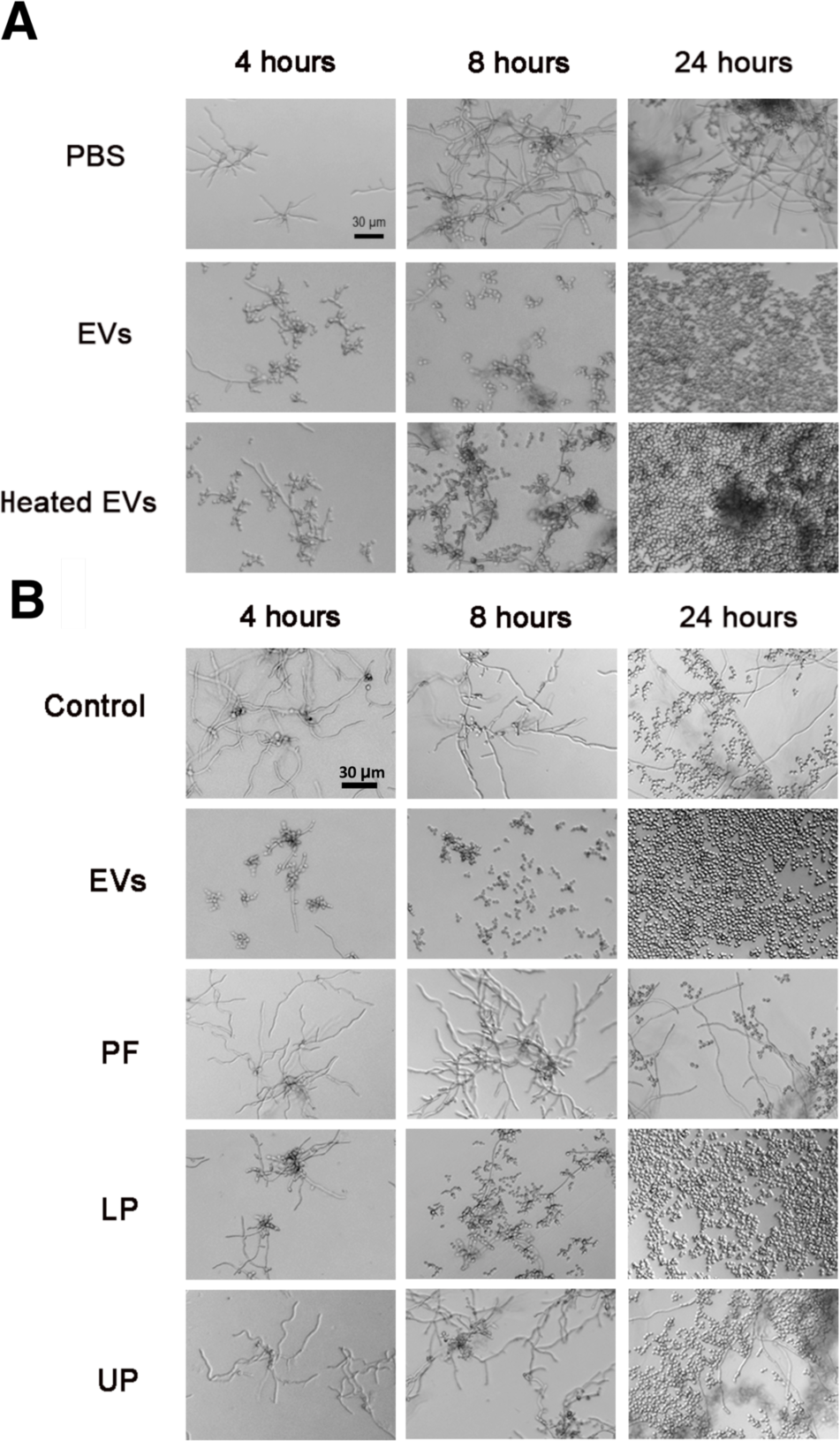
Thermo-resistant molecules carried by *Ca* EVs are associated with yeast-to-hyphae inhibitory effect. (A) Yeasts of *C. albicans* (90028) were inoculated into M199 (pH 7) in the presence or absence of *Ca* EVs or heat-treated EVs in concentration equivalent to 5 μg sterol/ml. Effect was visualized after 4, 8, and 24 h. PBS was added alone as a control. Bar, 30 μm. The results shown here are representative of three independent experiments. (B) Yeasts of *C. albicans* (90028) were inoculated into M199 in the presence or absence of *Ca* EVs, EV-derived protein-rich fraction (PF), or lipid lower (LP) or upper phase (UP) in concentration equivalent to 5 μg sterol/ml. Effect was visualized after 4, 8, and 24 h. PBS was added alone as a control. Bar, 30 μm.

To investigate whether the inhibitory effect of *Ca* EVs was not due to co-isolated contaminants, we fractioned our bioactive samples using size-exclusion chromatography (SEC), in a Sepharose CL-4B column. A single peak (fractions 8-10) containing proteins and lipids was observed (**Fig. S1**). DLS analysis confirmed the presence of EVs in fraction 8, corresponding to the most concentrated sample with higher amounts of lipids and proteins, but not in the adjacent ones. Two populations were characterized with average sizes previously reported in our studies^30^, ranging from 50-120 and 240-350 nm, and possibly corresponding to exosomes and ectosomes/microparticles, respectively. The inhibitory activity of the lipids from each fraction was then evaluated. Fraction 8 was able to inhibit the yeast-to-hypha differentiation of *C. albicans* after six hours of incubation (**Fig. S1**). Fraction 10, where the protein and lipid concentration were significantly reduced comparing to fraction 8, displayed a behavior similar to control samples, with clear hypha formation. None of the fractions where lipids and proteins were not detected showed inhibitory activity (data not shown).

### EVs produced by other fungal species affect yeast-to-hypha differentiation in *C. albicans*

On the basis of the lipid nature of the bioactive molecules studied in our work and on the fact that farnesol is a QSM lipid that controls yeast-to-hypha differentiation in *C. albicans* ^16, 19^, we hypothesized that its presence in EVs from *C. albicans* could result in filamentation inhibition. To confirm this possibility, we asked whether EVs from *S. cerevisiae*, which synthesizes and exports farnesol in very low amounts^50^, regulate morphogenesis in *C. albicans*. We also included EVs from *H. capsulatum* in these analyses, since farnesol has not been characterized as a lipid component in this fungus. We first investigated whether the strains used in our study produced detectable amounts of farnesol into the LP by reverse-phase thin-layer chromatography (RP-TLC or TLC). Whole cells were used in order to obtain larger amounts of sample (**Fig. S2**). The TLC showed a sharp band in the LP of *C. albicans* (*Ca*) with identical color and retention factor (R_f_) obtained for a farnesol (FOH) standard, strongly indicating the presence of this molecule in *C. albicans*. A band with similar R_f_ was observed in the LP of *H. capsulatum* (*Hc*); however, a distinct yellowish color was visualized. Thus, a more detailed structural analysis would be necessary to confirm the true nature of that band. Finally, no molecule migrating as farnesol was detected in the LP from *S. cerevisiae* (*Sc*) by TLC (**Fig. S2**). Nevertheless, EVs from both species were able to negatively impact *C. albicans* hyphal formation (**Fig. S3**), but the kinetics of inhibition (comparing 8 h vs. 24 h of incubation time) was considerably slower when compared to *Ca* EVs. With *Hc* EVs, the inhibitory effect was observed only after an overnight incubation. The first signals of inhibitory effect from *Sc* EVs were detected after 8 h of incubation. To confirm that this was not a pleiotropic effect caused by EVs produced by any of the organisms studied, we also tested the activity of EVs from murine macrophages-like (MO). Contrasting with the activity of fungal EVs, MO EVs did not interfere with yeast-to-hypha differentiation *in vitro* (**Fig. S3**). The fact that EVs from both *S. cerevisiae* and *H. capsulatum* inhibited *C. albicans* differentiation, even at a lower efficacy, suggests that EV molecules other than farnesol participate in the inhibitory mechanism. We next investigated whether other lipids could be potentially involved in the inhibitory effect of EVs.

### Terpenes are potentially the major compounds involved with *C. albicans* yeast-to-hypha inhibition

To investigate whether other lipids produced by *C. albicans* could also mediate the inhibitory activity of *Ca* EVs, we tested the different TLC bands from the *C. albicans* LP. Using a preparative TLC plate and staining the lipids with mild iodine vapor, to avoid lipid degradation, we obtained five (A-E) major fractions corresponding to separate bands (**Fig. S4)**. The ability of each fraction to inhibit yeast-to-hypha differentiation was investigated and only the one (band A) with an R_f_ similar to an authentic farnesol standard showed an active effect (**Fig. S4**). We then concentrated our efforts in the analysis of farnesol in fungal EVs. To that end, we followed a protocol previously used for farnesol extraction ^51^, and employed a highly sensitive GC-MS with selected-reaction monitoring (GC-MS/SRM) method, using an external *bona fide* standard of farnesol. As observed in Fig. 5A,B (black line), the farnesol standard (at 4.5 picomole/µL, 1 μL injection), is comprised of a mixture of four isomers (2-*cis*,6-*cis*-farnesol [Z,Z-farnesol], 2-*cis*,6-*trans*-farnesol [Z,E-farnesol], 2-*trans*,6-*cis*-farnesol [E,Z-farnesol], and 2-*trans*,6-*trans*-farnesol [E,E-farnesol]), which clearly separated from each other using the GC gradient conditions employed. The isomer profile was comparable to that previously reported ^52^. Under the experimental conditions used for the standard mixture, we could detect peaks (at 9.65 and 9.81 min) with SRM transitions (not shown) and full scan chromatogram elution time similar to Z,E-farnesol and E,E-farnesol isomers (peaks 3 and 5, respectively) in the 90028 yeast *Ca* EV sample (Fig. 5A,B). Table 1 shows the relative peak intensity of these farnesol isomers and other identified compounds found in *Ca* EVs and other lipid fractions. Beside the two farnesol isomers, the GC-MS analysis revealed that *Ca* EVs from contain high amounts of other terpenoids or isoprenoids such as geranylgeraniol isomers, and to a much lesser extent, squalene, which have not been previously reported in *C. albicans* (Table 1). The farnesol derivative 2,3-dihydro-6-*trans*-farnesol (6, 10-dodecadienol-1-ol, 3,7,11 trimethyl) (peak 1) and related isoprenoid, 2,3-dihydro-6-*trans*-geraniol (6,10,14-hexadecatrien-1-ol 3,7,11,15 tetramethyl) (peak 9), corresponding to farnesol and geranylgeraniol units lacking a double bond at C-2, respectively, were also characterized as major peaks in *Ca* EVs (Fig. 5, Tables 1 and 2, and **Fig. S7**). Of those isoprenoids, only 2,3-dihydro-6-*trans*-geraniol was also found in *Hc* and *Sc* EVs. None of the farnesol isomers or its derivative (2,3-dihydro-6-*trans*-farnesol) were detected in the EVs from *H. capsulatum* and *S. cerevisiae*, confirming that these molecules are only enriched in *Ca* EVs (Fig. 5, Tables 1 and 2**, and Fig. S7**). Five fatty acids were also identified as major *Ca* EV components, including octadecanoic (stearic acid, C18:0) and octadecenoic (oleic acid, C18:1) acids, and to a much lesser extent, (9Z)-hexadec-9-enoic (palmitoleic acid, C16:2), dodecanoic (lauric acid, C12:0), and tetradecanoic (myristic acid, C14:0) acids (Fig. 5 and Table 1). Fatty acids were also found in EVs from *H. capsulatum* (C12:0, C14:0, and C18:0) and *S. cerevisiae* (C12:0, C14:0, C16:1, C18:0, and C18:1) (Fig. 5 and Table 1). Ascorbic acid 2,6-dihexadecanoate was found in EVs from the three fungal species (Fig. 5 and Table 1). This compound had no yeast-to-hypha inhibitory effect (data not shown). We also characterized the major lipids found in the LP from *C. albicans* and its bioactive TLC-derived band (**Fig. S4A, B**). Equivalent amounts of 2,3-dihydro-6-*trans*-farnesol and E,E-farnesol were detected in the *C. albicans* LP (Fig. 5). 2,3-dihydro-6-*trans*-geraniol was also identified in *Ca* LP (Tables 1 and 2). Moreover, the free fatty acids C12:0, C14:0, C18:0, and C18:1 were also identified in this fraction. Finally, we resolved the lipids from *C. albicans* EVs by RP-TLC and the band corresponding to farnesol R_f_ was also analyzed. We found a high similarity between the total EV lipids and this band, with a clear enrichment of farnesol isomers and farnesol-like structures, and other isoprenoids/terpenoids (Tables 1 and 2). In addition, we observed an enrichment of the fatty acid C12:0 and dodecanoic acid, undecyl ester (C23:0-O2) (Fig. 5A and Table 1). Using the standard calibration curve from the d6-E,E-farnesol isomer (**Fig. S5**), we were able to quantify the amounts of each isoprenoid/terpenoid found in fungal EVs (Table 2), confirming the results above and a significant enrichment of isoprenoids in EVs carried by *C. albicans*.

**Figure 5.**
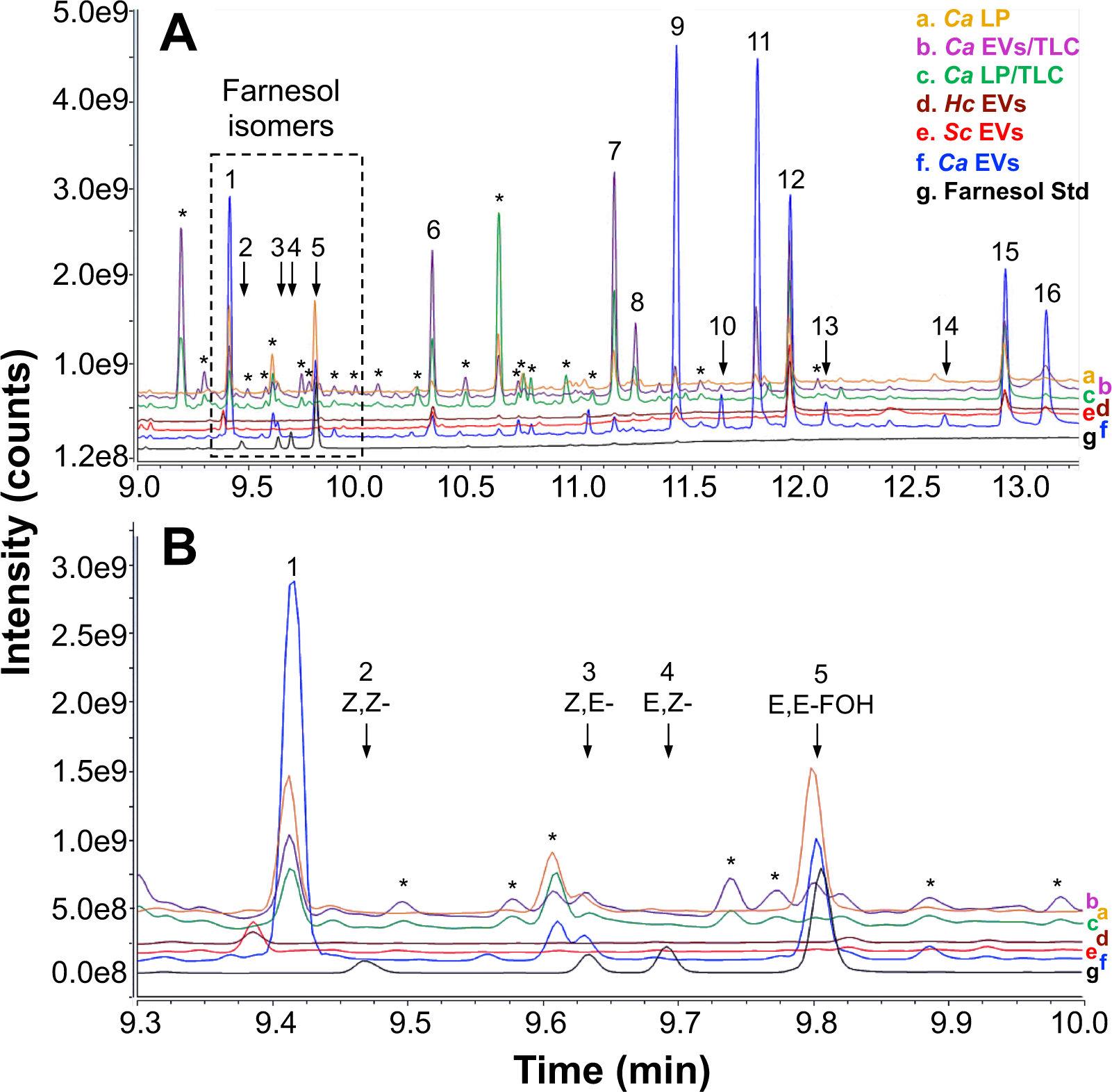
GC-MS analysis of fungal lipids and EVs. Lipids were extracted from *Ca* LP fraction and EVs, and from RP-TLC band co-migrating with farnesol in those two samples. Lipids were also extracted from *Hc* and *Sc* EVs. Resulting samples were analyzed by GC-MS/MS. All (A) GC-MS full scan chromatogram. The chromatogram region containing the four farnesol isomers is indicated (dashed rectangle) and shown in detail in (B). A standard mixture containing the four farnesol isomers (Z,Z-, Z,E-, E,Z-, and E,E-FOH) was used as reference (black line/trace). The GC-MS full-scan traces were overlaid and labelled (**a-g**) to facilitate visualization and comparison. Farnesol isomers and peaks of interest are indicated numerically and annotated in Table 1. Asterisks indicate contaminants (e.g., phthalates and other plasticizers) or compounds not identified in the library (low structural match probability).

**Table 1.** Major lipid compounds identified by GC-MS/MS in the lipid LP fractions and EVs of *C. albicans*, and EVs from *H. capsulatum*, and *S. cerevisiae*.

| Peak number | RT (min) | Compound | Fungal Preparation/Fraction |  |  |  |  |  |
| --- | --- | --- | --- | --- | --- | --- | --- | --- |
|  |  |  | *C. albicans* |  |  |  | *H. capsulatum* | *S. cerevisiae* |
|  |  |  | LP | EVs | LP/TLC | EVs/TLC | EVs | EVs |
|  |  |  | Peak Relative Intensity (%)<sup>a</sup> |  |  |  |  |  |
| 1 | 9.42 | 2,3-dihydro-6-*trans*-farnesol (6,10-dodecadien-1-ol, 3,7,11 trimethyl) | 93 | 63 | 32 | 25 | nd | nd |
| 2 | 9.47 | 2,6-*cis*-farnesol (Z,Z-farnesol) | nd | nd | nd | nd | nd | nd |
| 3 | 9.63 | 2-*cis*,6-*trans*-farnesol (Z,E-farnesol) | nd | 4 | nd | 7 | nd | nd |
| 4 | 9.68 | 2-*trans*,6-*cis*-farnesol (E,Z-farnesol) | nd | nd | nd | nd | nd | nd |
| 5 | 9.81 | 2,6-*trans*-farnesol (E,E-farnesol) | **100** | 20 | 5 | 9 | nd | nd |
| 6 | 10.33 | dodecanoic acid (lauric acid, C12:0) | 14 | 5 | 57 | 67 | 25 | 23 |
| 7 | 11.15 | tetradecanoic acid (myristic acid, C14:0) | 41 | 4 | 97 | **100** | 6 | 4 |
| 8 | 11.25 | dodecanoic acid, undecyl ester (C23:0-O2) | nd | nd | 30 | 32 | 6 | 4 |
| 9 | 11.42 | 2,3-dihydro-6-*trans*-geraniol (6,10,14-hexadecatien-1-ol 3,7,11,15 tetramethyl) | 14 | **100** | 8 | 12 | 19 | 9 |
| 10 | 11.63 | geranylgeraniol (isomer 1)<sup>c</sup> | nd | 9 | nd | 3 | nd | nd |
| 11 | 11.79 | geranylgeraniol (isomer 2)<sup>d</sup> | 10 | 97 | nd | 39 | nd | nd |
| 12 | 11.93 | ascorbic acid 2,6-dihexadecanoate | 72 | 60 | **100** | 68 | **100** | **100** |
| 13 | 12.11 | (9Z)-hexadec-9-enoic acid (palmitoleic acid, C16:2) | nd | 7 | 3 | 3 | nd | 4 |
| 14 | 12.64 | squalene | nd | 3 | nd | nd | nd | nd |
| 15 | 12.92 | octadecanoic acid (stearic acid, C18:0) | 41 | 41 | 62 | 30 | 38 | 36 |
| 16 | 13.12 | octadecenoic acid (oleic acid, C18:1) | 3 | 30 | 5 | 10 | nd | 14 |

<sup>a</sup> Not detected (below the method's limit of detection).
<sup>b,c</sup> We were unable to determine the precise structures of these geranylgeraniol geometric isomers through fragmentation spectral library matching.

**Table 2.** Quantification of farnesol and farnesol-like structures in lipid LP fractions and EVs of *C. albicans*, and EVs from *H. capsulatum* and *S. cerevisiae*.

| Fungal fraction/preparation | Farnesol isomers (ng/100 µg total lipid or protein)<sup>a</sup> |  |  | Farnesol-like structures (ng/100 µg total lipid or protein)<sup>a</sup> |  |  |
| --- | --- | --- | --- | --- | --- | --- |
|  | 2,3-Dihydro-6-*trans*-farnesol | Z,E-FOH | E,E-FOH | 2,3-dihydro-6-*trans*-geraniol | Geranylgeraniol (isomer 1) | Geranylgeraniol (isomer 2) |
| *Ca* LP | 16,3 | 1,8 | 19 | 2,1 | nd | 1,2 |
| *Ca* LP/TLC | 2,1 | nd<sup>b</sup> | nd | nd | nd | nd |
| *Ca* EVs | 33,7 | 1,6 | 9,7 | 67 | 4,1 | 63,4 |
| *Ca* EVs/TLC | 48,8 | 7,5 | 9,6 | 21 | nd | 77,4 |
| *Hc* EVs | nd | nd | nd | 4,7 | nd | 1,9 |
| *Sc* EVs | nd | nd | nd | 1,2 | nd | nd |

<sup>a</sup> LP and LP/TLC fractions were normalized by 100 µg of total lipid, whereas all EV preparations were normalized by 100 µg of protein.
<sup>b</sup> Not detected (below the method's limit of detection).

### EVs from *C. albicans* yeast cells reverse the yeast-to-hypha differentiation and their inhibitory effect is not reversed by db-cAMP

Given the presence of additional compounds in *Ca* EVs that could potentiate their effects in morphogenesis we next investigated whether *Ca* EVs are able to reverse the yeast-to-hypha differentiation induced by M199 medium. Yeast cells were plated onto M199 medium supplemented with tyrosol for stimulation of filamentation and hyphal formation was observed after 4 h of induction (**Fig. S6**), at which time *Ca* EVs were added to the wells. Yeast cells were visualized 4 h after the addition of *Ca* EVs to the filamentous cultures, indicating that fungal EVs are able to efficiently reverse the yeast-to-hypha morphogenesis even after the highly regulated process had been initiated. As previously demonstrated ^53^, addition of farnesol to the wells did not reverse this effect (data not shown). Farnesol stops *C. albicans* filamentation by inhibiting adenylate cyclase (Cyr1) ^54^. This effect is rescued *in vitro* by the addition of exogenous db-cAMP ^55^ and it was also confirmed in our experiments (data not shown). Addition of db-cAMP to yeasts pre-treated with EVs did not rescue the filamentation, suggesting that the inhibitory effect was not dependent on Cyr1 activity (data not shown).

### Transcriptomic alteration of *C. albicans* in response to *Ca* EV treatment

We examined the changes in *C. albicans* gene expression under filament-repressing conditions as a result of *Ca* EV treatment. The cells were grown in the presence of three filament-repressing agents, the *Ca* EVs and LP fraction, and standard farnesol. Control cells were maintained as yeast forms at pH 4 or as filaments at pH 7. In average, we obtained at least 28 times of genome coverage and more than 90% of the reads mapped to the *C. albicans* genome (**Suppl. Table 1**). We next assessed if the treatment with the filament-repressing agents would lead to similar alterations in the cell transcriptome by applying a principal component analysis (PCA) (Fig. 6A). Each condition clustered separately, indicating distinct responses. The replicates clustered together, suggesting a high level of correlation between them and reinforcing data reliability and the reproducibility of our results (Fig. 6A). When compared to the control, a total of 411 mRNAs were differentially expressed in the samples treated with farnesol, 334 with the LP and 304 with the *Ca* EVs (Fig. 6B). The complete differential expression gene analysis (DEG) data are available (**Supp. Tables 2-5**). In addition, 100 transcripts were common to all three conditions studied (Fig. 6B). Analysis of the transcripts altered by the EV treatment showed upregulation of 143 transcripts and 161 downregulated genes (Fig. 6C and D). A protein interaction network analysis revealed a clear association between the transcripts, as concluded from the clusters formed by the STRING (Search Tool for the Retrieval of Interacting Genes) tool (Fig. 6E). An enrichment in DNA metabolism was evident (p-value < 1.0e^-16^). Gene ontology analysis identified DNA replication, mismatch repair and homologous recombination as key events (Table 3). Fourteen transcripts were linked to DNA replication (Fig. 6F). This result agrees with the observation that EV treatment increased yeast growth rates (Fig. 1A). As for the comparison between the transcripts affected the three filament-repressing agents, the effects induced by LP and EV were more mutually related than each of them was related to the effects induced by farnesol (Table 3). In the set of downregulated transcripts after, most of the affected transcripts were related to adhesion and extracellular proteins (Table 4).

**Figure 6.**
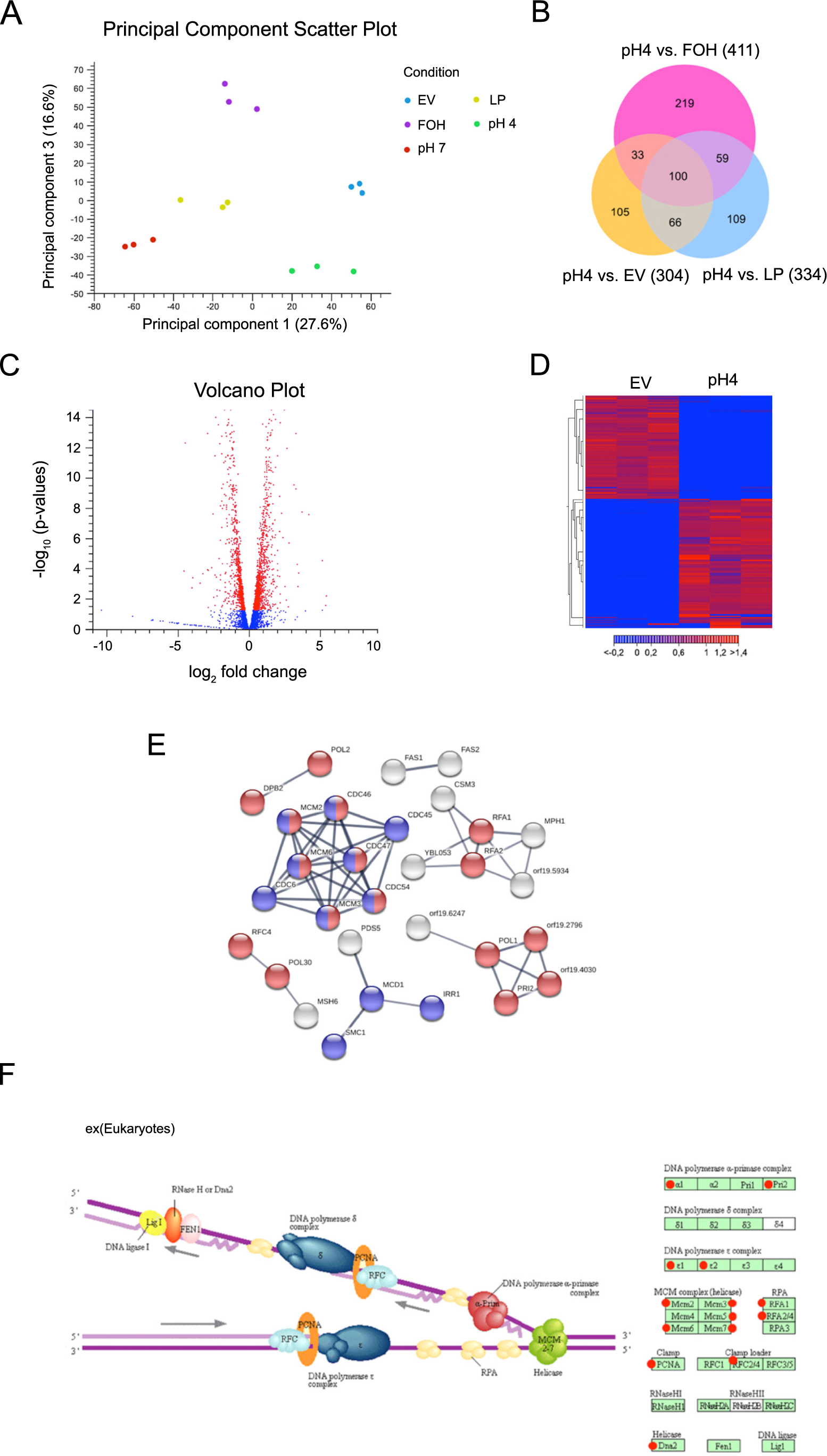
Transcriptomic alterations of *C. albicans* in response to EV treatment. (A) PCA plot showing the level of correlation and reproducibility among samples: blue (*Ca* EVs), purple (farnesol, FOH), yellow (*Ca* LP) and the controls: red (hyphae – pH 7) and green (yeast – pH 4). (B) Venn diagram showing the transcripts that are common to and exclusive to the conditions analyzed. Only transcripts that passed the statistical filter were selected for the analysis (log fold change ³ 3 £ −3 and FDR adjusted p-value < 0.05). All the conditions were compared to the yeast cell control (pH4). (C) Volcano plot showing the significantly upregulated (left) and downregulated (right) genes during the EV treatment. The red dots are all the transcripts that passed the cutoff of log fold change 3 and FDR of 0.05. (D) Heat map of gene expression data during the EV treatment. The color indicates the degree of correlation (red, high correlation; blue, low correlation; the scale at the bottom is represented in log scale). (E) Using the STRING tool it is possible to observe the enrichment for DNA replication (FDR adjusted p-value 8.4e-14) are indicated as red dots, while the cell cycle (FDR adjusted p-value 4.15e-6) are indicated as blue dots, and the gray dots are proteins associated to other functions. The thickness of the line indicates the degree of confidence prediction of the interaction, based only in experimentally validated data. (F) KEGG analysis indicated that a total of 14 genes out of 30 involved in DNA replication pathway are indicated as red dots.

**Table 3.** Kegg Pathway enrichment - transcripts upregulated following *Ca* EV treatment.

| #term ID | term description | observed gene count | background gene count | FDR |
| --- | --- | --- | --- | --- |
| cal03030 | DNA replication | 16 | 32 | 0.00% |
| cal00240 | Pyrimidine metabolism | 12 | 67 | 0.00% |
| cal03430 | Mismatch repair | 8 | 21 | 0.00% |
| cal03410 | Base excision repair | 6 | 17 | 0.01% |
| cal03440 | Homologous recombination | 6 | 17 | 0.01% |

**Kegg Pathway enrichment - transcripts Upregulated during the LP (Lipidic Lower Phase) treatment**
| #term ID | term description | observed gene count | background gene count | FDR |
| --- | --- | --- | --- | --- |
| cal03030 | DNA replication | 12 | 32 | 0.00% |
| cal04111 | Cell cycle - yeast | 14 | 85 | 0.00% |
| cal04113 | Meiosis - yeast | 11 | 71 | 0.01% |
| cal00220 | Arginine biosynthesis | 5 | 16 | 0.16% |
| cal00240 | Pyrimidine metabolism | 8 | 67 | 0.48% |

**Kegg Pathway enrichment - transcripts Upregulated during the Farnesol (FOH) treatment**
| #term ID | term description | observed gene count | background gene count | FDR |
| --- | --- | --- | --- | --- |
| cal01100 | Metabolic pathways | 50 | 651 | 0.01% |
| cal01110 | Biosynthesis of secondary metabolites | 25 | 270 | 0.22% |
| cal00920 | Sulfur metabolism | 5 | 13 | 0.80% |
| cal01230 | Biosynthesis of amino acids | 12 | 99 | 1.15% |
| cal01130 | Biosynthesis of antibiotics | 18 | 200 | 1.31% |

**Table 4.**
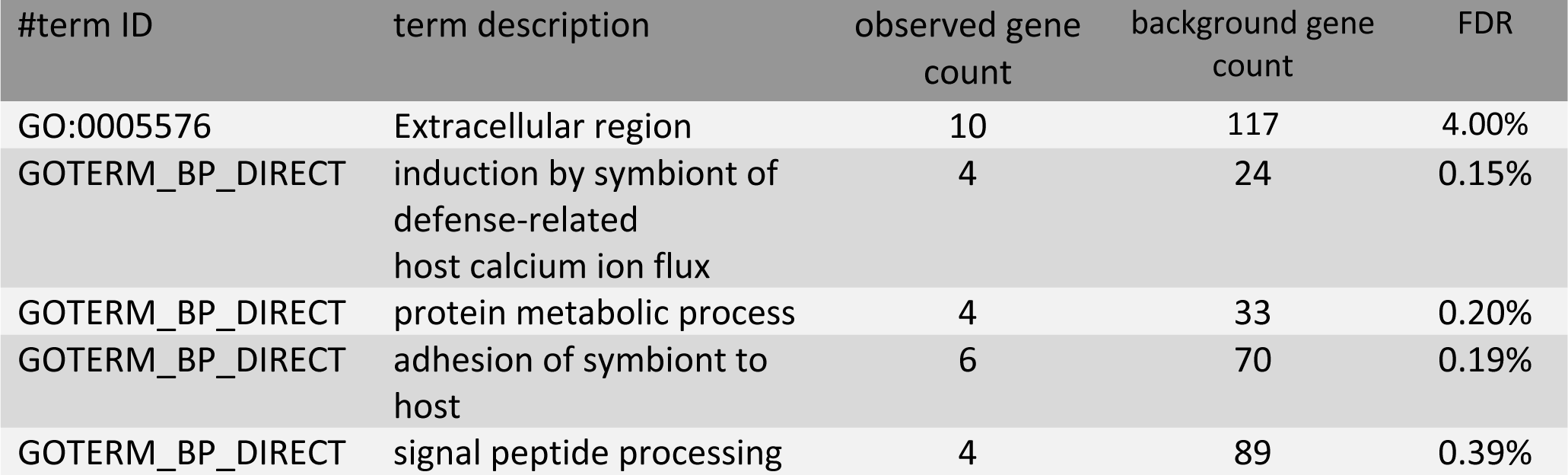
Kegg Pathway enrichment – transcripts downregulated following Ca EV treatment.

| #term ID | term description | observed gene count | background gene count | FDR |
| --- | --- | --- | --- | --- |
| GO:0005576 | Extracellular region | 10 | 117 | 4.00% |
| GOTERM_BP_DIRECT | induction by symbiont of defense-related host calcium ion flux | 4 | 24 | 0.15% |
| GOTERM_BP_DIRECT | protein metabolic process | 4 | 33 | 0.20% |
| GOTERM_BP_DIRECT | adhesion of symbiont to host | 6 | 70 | 0.19% |
| GOTERM_BP_DIRECT | signal peptide processing | 4 | 89 | 0.39% |

### Treatment with EVs reduces ability of *C. albicans* to invade solid medium and decreases fungal virulence in a *Galleria mellonella* model of candidiasis

Due to the well-established connections between pathogenesis and morphological transitions in *C. albicans* ^8^, we investigated whether treatment with *Ca* EVs could abrogate the ability of the fungus to invade solid medium, a consequence of filamentous growth regularly correlated with virulence ^56^. Colonies of *C. albicans* cultivated in M199 agar medium produced normal hypha at the edges after 5-6 days of invasive growth (Fig. 7A**, upper left panel**). In contrast, after 5-6 days no hyphae were visualized when yeast cells were pre-treated with *Ca* EVs (Fig. 7A**, upper right panel**). In fact, under *Ca* EV-treatment conditions, hyphal formation was only observed after 10-11 days of incubation (data not shown). These results were confirmed using the plate-washing method ^56^. After 10 days of growth the colonies (control and EV-treated) were washed using the same flow rate, water temperature and period of time (Fig. 7A**, lower panels**), producing results that were similar to those obtained under standard conditions.

**Figure 7.**
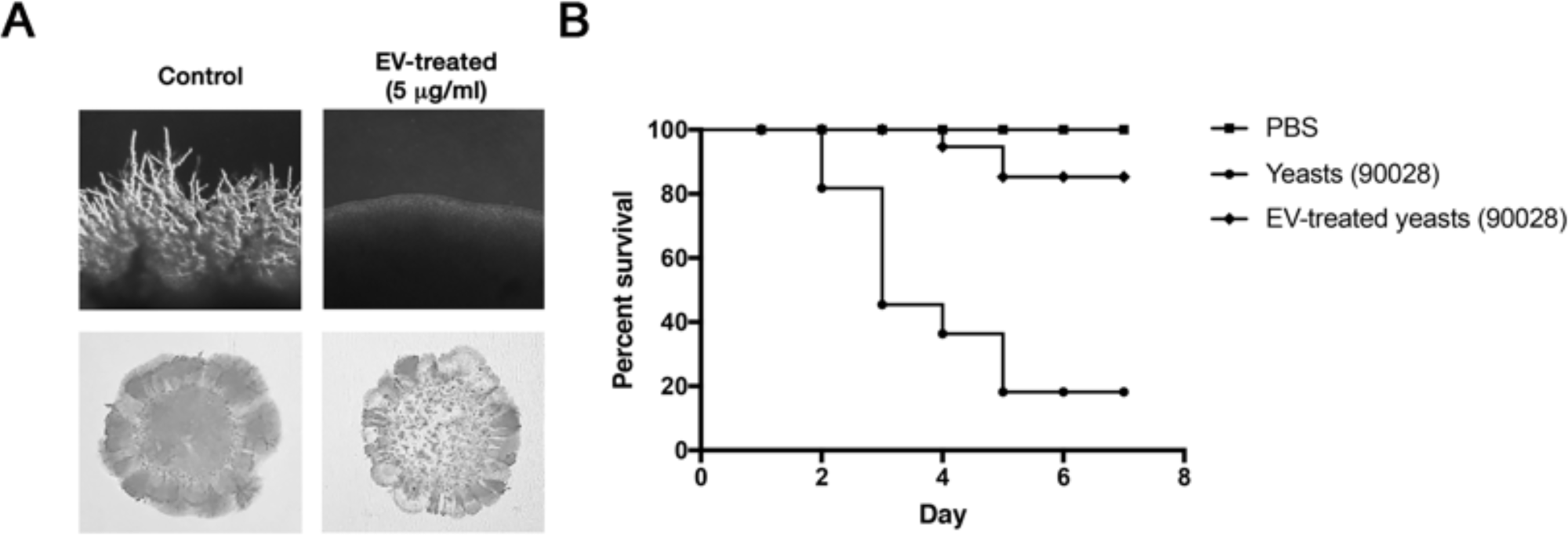
*Ca* EVs impact invasive assay and virulence in *G. mellonella*. (A) Yeasts of *C. albicans* (90028) were treated with PBS (control) or *Ca* EVs in PBS (5 μg/ml) and then plated onto M199 agar for 7 days. Presence of filamentation (upper panels) and agar invasion (lower panel) was observed only under control conditions. (B) *Galleria mellonella* larvae were infected with 10^5^ yeasts of *C. albicans* untreated or pretreated with *Ca* EVs (5 μg/ml – 24 h) and the number of dead larvae was monitored daily. PBS indicates control larvae injected with PBS. Experiments were performed in biological duplicate with similar results.

The capacity to reduce the adhesive and invasive abilities of *C. albicans* indicated that pre-incubation of yeast cells with *Ca* EVs could decrease the virulence of *C. albicans*. To investigate whether the treatment of yeast cells with EVs would impact pathogenesis we used *G. mellonella* as a model. Infection of *G. mellonella* larvae with a lethal inoculum of yeast cells of *C. albicans* cultivated in standard medium without *Ca* EVs caused melanization and death in 80% of the animals in less than one week (Fig. 7B). Remarkably, when the pre-treated yeast cells were used to infect the larvae only 10% of the animals died (Fig. 7B). We then evaluated the ability of EVs to impair differentiation in the presence of serum, a major hyphae-inducing factor in mammalian hosts^57^. Our results demonstrated that *C. albicans* EVs are able to inhibit yeast-to-hypha differentiation in the presence of serum and CO_2_ (data not shown). Together, these data suggested that *C. albicans* EVs could directly affect the pathogenesis of *C. albicans* during the infection.

## Supplementary material (uploaded)

**Supplemental table 1:** RNA-seq mapping statistics

**Supplemental table 2:** Differential gene expression analysis of the transcriptomic data from the yeast cells treated with the Extracellular vesicles (EV).

**Supplemental table 3:** Differential gene expression analysis of the transcriptomic data from the hypha cells.

**Supplemental table 4:** Differential gene expression analysis of the transcriptomic data from the yeast cells treated with farnesol (FOH).

**Supplemental table 5:** Differential gene expression analysis of the transcriptomic data from the yeast cells treated with the Lipidic lower phase (LP).

**Supplemental Figure 1:** Size-exclusion chromatography (SEC) and DLS of EVs from *C. albicans.* The *Ca* EVs released into medium was enriched by SEC, using Sepharose CL-4B, and the 500-μL fractions collected as described in Methods. Each fraction was quantified according to its protein and sterol content. Fraction 8 displayed two populations of EVs with effective diameters usually found in *Ca* EVs. Yeasts of *C. albicans* (90028) were inoculated into M199 in the presence of each fraction to evaluate their yeast-to-hypha inhibitory activity. Fractions containing EVs were normalized according to the sterol content and 5 μg/ml were used. Fractions with no protein or sterol were also tested and 80% of the total content was used. None of these fractions showed any inhibitory activity. Effect was visualized after 6 and 24 hours. PBS was added alone as a control. Bar, 30 μm.

**Supplemental Figure 2:** Major lipids from *C. albicans, H. capsulatum* and *S. Cerevisiae* resolved by TLC. LP enriched in non-polar lipids from each fungal species was obtained as described in Methods. *C. albicans* (C*a*), *H. capsulatum* (*Hc*), and *S. cerevisiae* (*Sc*) lipid samples were resolved by reverse-phase TLC using acetone as solvent system and stained with ferric chloride-sulfuric acid reagent. Farnesol (FOH), indicated by an arrow, was used as standard.

**Supplemental Figure 3:** Effect of EVs from different fungal species and macrophages over yeast-to-hypha differentiation. Yeasts of *C. albicans* (90028) were inoculated into M199 in the presence or absence of EVs from *H. capsulatum* (*Hc*), *S. cerevisiae* (*Sc*), and macrophage-like RAW cell line (MO) at the equivalent concentration of 5 μg sterol/ml. Effect was visualized after 4, 8, and 24 h. PBS was added alone as a control (CT). Bar, 30 μm.

**Supplemental Figure 4:** Effects of bands isolated by preparative reverse-phase thin layer chromatography (RP-TLC). (A) Lipids from LP of *C. albicans* strain 90028 (*Ca* LP) were resolved by RP-TLC and the bands (a-e) stained with I_2_ vapor were scraped and eluted from the silica (see Methods). Farnesol (FOH), indicated by an arrow, was used as standard. (B) Yeasts of *C. albicans* (90028) were inoculated into M199 in the presence or absence of each eluted RP-TLC band (suspended in DMSO as described in Methods) Effect was visualized after 6 and 24 h. PBS-DMSO was added alone as a control. Bar, 30 μm.

**Supplemental Figure 5:** Standard calibration curve of d6-farnesol E,E isomer. d6-Farnesol E,E isomer (Avanti Polar) was diluted to concentrations of 0.1, 1.0, 2.5, and 5.0 ng/µL. Peak areas were imported to GraphPad v9.0.0 for graph and linear regression generation. The 95% confidence interval (discontinued line) of the curve is indicated.

**Supplemental Figure 6:** *C. albicans* EVs reversed the yeast-to-hypha differentiation. Yeasts of *C. albicans* (90028) were inoculated into M199 and incubated for 4 hours to induce hyphae formation. Then PBS (control), *Ca* EVs (5 μg/ml) or a combination of EVs (5 μg/ml) and tyrosol (200 μM) were added to the wells. Cell morphology was visualized after 8, 12 and 24 hours. Bar 30 μm.

**Supplemental Figure 7:** Major terpenoids or isoprenoids identified in *C. albicans* EVs and lipid extracts.

**Supplemental video 1:** *C. albicans* growth under inducing filamentous conditions. Yeasts of *C. albicans* (90028) were inoculated into M199 medium to induce filamentation. Cellular density was monitored up to 800 min. This video is representative of two independent experiments with similar results.

**Supplemental video 2:** *C. albicans* growth under inducing filamentous conditions in the presence of EVs. Yeasts of *C. albicans* (90028) were inoculated into M199 medium to induce filamentation in the presence of EVs (5 μg/ml based on sterol content). Cellular density was monitored up to 800 min. This video is representative of two independent experiments with similar results.

## Discussion

*C. albicans* morphogenesis has been associated with fungal colonization and adaptation to distinct micro environmental conditions ^5, 6^. Additionally, morphological transition is essential during biofilm formation and disease development ^8, 9, 58, 59^. In fact, inhibition of morphological switching has been considered a therapeutic target for candidiasis ^59^. Environmental and auto regulatory molecules involved with morphogenesis in *C. albicans* have been extensively investigated ^16, 18–21^. However, the knowledge on *C. albicans* molecules inducing specific morphological stages is very limited, as are the mechanisms by which these compounds are released to the extracellular environment. Farnesol and tyrosol are examples of QSM produced by *C. albicans* that regulate morphogenesis and biofilm formation ^16, 18, 48, 60^. Considering that fungal EVs have been associated with cell-cell communication and biofilm matrix production ^28, 44^, we hypothesized that these membranous compartments could be involved with biofilm formation and morphogenesis of *C. albicans.* Our results demonstrate that EVs released by fungi are able to directly affect yeast growth and inhibit biofilm formation, through blocking of yeast-to-hypha differentiation and stimulation of hypha-to-yeast differentiation.

The impact of EVs produced by *C. albicans* strain 90028 (*Ca* EVs) was investigated during growth and biofilm formation. Remarkably, addition of *Ca* EVs to *C. albicans* yeast cells culture induces faster growth. In addition, our results suggest that *Ca* EVs partially mimic the effect of farnesol, displaying a filamenting-suppressing activity and inhibiting biofilm formation, which could also be observed in our transcriptomic analysis. However, this effect was prolonged in comparison to the activity of the commercial farnesol, indicating that yeast-to-hyphae morphogenesis was temporarily turned off. Commercial farnesol mixtures usually contain its four isomers (predominantly, the E,E-farnesol) and has been extensively used to investigate as the QSM molecule produced by *C. albicans*. The exact mechanism and signaling pathways triggered by *Ca* EVs that promote the long lasting effect were not investigated, but it could involve epigenetic changes as reported previously for the white-opaque transition^61^. Additional changes are also expected after *Ca* EV treatment and must be further investigated. For instance, *Ca* EV-treated cells lost their ability to adhere to the plate’s wells after 7 h of incubation suggesting that treatment with *Ca* EVs changed surface composition. Although the activity of *Ca* EVs was dose dependent, the inhibitory effect was apparently not linked to yeast density in fungal cultures. These data suggest that the *Ca* EV components involved with morphogenesis must reach a threshold concentration to inhibit differentiation, a circumstance that is well-known for the QSM farnesol and tyrosol ^18, 48^. This hypothesis is in agreement with the ordinary, physiological transition between yeast and hyphal forms, which would not occur regularly if EVs or other extracellular regulators were inhibitory at minimum concentrations. Altogether, these results indicate that *Ca* EVs carry structures able to regulate growth and morphogenesis in *C. albicans*.

In order to confirm that *C. albicans* uses EV export as a general mechanism to control yeast growth and morphogenesis *in vitro,* we isolated *Ca* EVs from two other strains with distinct filamentation abilities and virulence profiles and tested their ability to inhibit yeast-to-hyphae differentiation. Our data revealed that three different strains of *C. albicans* produce EVs with similar properties, which were also comparable to fungal EVs, recently characterized by Reis and colleagues using similar detection approaches with *C. gattii* ^46^. In addition, small differences in size were observed when the distinct strains were compared, confirming previous data published by our group showing that *C. albicans* strains can produce EVs with distinct sizes ^30^. Morphological differences of EVs are usually attributed to changes in composition and / or biogenesis pathways. The differentiation process of all strains was affected by the *Ca* EVs. In addition, *Ca* EVs from the different strains were able to promote the inhibitory effect. Thus, the *Ca* EV capacity of inhibiting yeast-to-hypha differentiation seems to be conserved in the three strains used in our study. On the other hand, the ability of EVs isolated from different species, such as *H. capsulatum* and *S. cerevisiae,* to inhibit *C. albicans* filamentation was considerably reduced, suggesting that EVs from *C. albicans* should have a distinct cargo that influences morphogenesis.

A number of structures carried by fungal EVs could be involved with growth and morphogenesis control, including lipids, proteins and RNA. The participation of thermo-labile molecules, including a number of proteins, was discarded since the filamentation inhibition was observed even when the *Ca* EVs were heated. Since lipids are thermostable, these results suggested that a lipid compound could be responsible by the *Ca* EV-mediated inhibitory activity. Given that farnesol is a lipid that regulates morphogenesis in *C. albicans,* we speculated that this molecule could be the *Ca* EV components mediating the yeast-to-hypha inhibitory effect. Our TLC results confirmed the presence of a farnesol band with inhibitory activity exclusively in *C. albicans* lipids. However, we could not rule out that the colorimetric method used in our experiments was not sensitive to detect low amounts of farnesol in *H. capsulatum* and *S. cerevisiae* strains. In addition, it was still necessary to confirm that farnesol is addressed to *C. albicans* EVs at appropriate concentrations.

Our GC-MS results strongly indicated the presence of lipids in *C. albicans* EVs that could interfere with morphogenesis, including a family of terpenes and middle chain fatty acids^48, 62, 63^. Besides farnesol, other sesquiterpens and diterpenes never reported previously in *C. albicans* were the major compounds characterized in the *Ca* EVs. Of note, the proportion of the terpene species in *C. albicans* EVs was distinct from the LP that showed the highest amounts of farnesol isomers. In addition, diterpenes were found in very low amounts in EVs released by *S. cerevisiae* and *H. capsulatum* which could explain why they have a reduced activity to inhibit differentiation when compared to *Ca* EVs.

Our GC-MS data also showed the presence of different fatty acids species in EVs from *C. albicans.* Earlier studies demonstrated that long chain fatty acids had no effect on fungal differentiation, including octadecanoic an octadecenoic acids, major components in *C. albicans* EVs^64^. However, they also showed that butyric acid was able to inhibit *C. albicans* germination. Willems and colleagues^62^ showed that decanoic acid inhibits *C. albicans* filamentation without affecting fungal growth. In addition, Lee and colleagues^63^ demonstrated that C7:0, C8:0, C9:0, C10:0, C11:0 and C12:0 impacted biofilm formation and filamentation. Furthermore, C7:0 and C9:0 also modified the expression of biofilm and hyphal grow regulating genes. Finally, the pre-treatment of *C. albicans* with C9:0 reduced the virulence of this fungus in a *C. elegans* model of candidiasis^63^. Thus, the presence of C10:0 and C12:0 in *C. albicans* EVs could directly contribute with filamentation inhibition.

Our results indicate that farnesol is not the only EV molecule regulating *C. albicans* morphogenesis. Our data demonstrate that *Ca* EVs reversed the hypha formation even under conditions where filamentation was already initiated and this reversion was not observed when farnesol was added to the cultures in previous studies^16, 50^. Remarkably, *Ca* EVs were able to reverse the filamentation even when induced by M199 supplemented with tyrosol. In addition, farnesol has a short time effect and is not able to modulate yeast growth positively^16^. In fact, higher concentrations of farnesol completely inhibited *C. albicans* growth ^65^. Furthermore, addition of db-cAMP did not rescue the inhibitory effect of *Ca* EVs, indicating that this inhibitory effect was independent from the adenylate cyclase activity and distinct from the regulation stimulated by farnesol. Finally, according to the GC-MS analysis the concentration of farnesol carried by *Ca* EVs seems to be below the limit required for its activity. The concentration of farnesol usually recommended to inhibit *C. albicans* differentiation is in the range of 1 to 150 μM^16, 21, 53, 55, 65^. According to our quantitative analysis the sterol:protein weight ratio is 1:20 (data not shown). Most of our experiments were developed using a sterol concentration of 5 μg/ml, corresponding to a protein concentration of 100 μg/ml of protein. Considering the lowest concentration of farnesol used in previous studies (1 μM), the amount of farnesol required for yeast-to-hyphae inhibition is 222.3 ng/mL. As demonstrated in Table 2, an EV suspension able to inhibit filamentation contains approximately 11.3 ng/ml of farnesol, corresponding to the sum of both Z,E-farnesol and E,E-farnesol isomers. In conclusion, *Ca* EVs carry a family of lipids, including diterpenes, sesquiterpenes and fatty acids that altogether can control the morphological changes in this organism. Future studies must be performed to evaluate the activity of each terpene species and the combination of these structures with middle chain fatty acids to inhibit *C. albicans* filamentation.

It has been speculated that the reduced size and round shape of yeast cells make them the putative forms used by *C. albicans* to disseminate in the host. However, this possibility is questionable due to the presence of serum in the blood, which would promptly induce germ tube and hypha formation^57^. Our experiments show that *Ca* EVs inhibit yeast-to-hypha transition even in the presence of FBS serum and 5% of CO_2,_ indicating that morphogenesis could be controlled by EVs during mammalian infections. This possibility is corroborated by the long-term effect manifested by *Ca* EVs *in vitro*. Thus, *Ca* EVs would make hypha-to-yeast differentiation feasible *in vivo*, which could possibly explain how this fungus modulates differentiation to disseminate.

It was possible to observe that the transcriptomic analysis of yeasts treated with *Ca* EVs triggered a specific response in gene expression, compared to the other treatments tested. The pathways upregulated were related to DNA metabolism, like the replication, mismatch repair and homologous recombination. This could be associated with the higher rates of cell growth observed when the cells were treated with *Ca* EVs. The EVs have been implied in cell cycle or cell growth on other organisms; for example, in cancer cells, EVs can promote tumor growth ^66, 67^. In tumor and stromal cells models, EVs are associated with cell proliferation^66^. In *S. cerevisiae*, EVs were recently associated with cell wall remodeling and enhancement of cell viability as the EV preparations increased the viability of yeast cells in the presence of caspofungin ^68^.

Based on the results discussed above, we hypothesized that pre-treatment of yeast cells *in vitro* with EVs would also decrease *C. albicans’* ability to invade host cells and cause disease. This possibility was reinforced by the fact that there was a delay during filamentation in the edge of colonies formed by *Ca* EV-treated yeast cells during invasive agar tests. To correlate the inhibitory effect of *Ca* EVs with the decrease in fungal invasion and virulence, we investigated the ability of pre-treated yeast cells to kill *G. mellonella* larvae. Indeed, *Ca* EV treatment significantly attenuated the ability of *C. albicans* to cause larvae death. Melanization was observed only in larvae infected with untreated yeasts, confirming stress due to candidiasis progress. This result indicates that by inhibiting yeast-to-hypha differentiation, *Ca* EVs decrease fungal virulence. In addition, it validates the idea that controlling filamentation would be a potential strategy as a therapeutic strategy^59^.

In our study, the transcripts downregulated after *Ca* EV treatment were mainly associated with adhesion to host cells. These results are possibly related to the hypovirulent phenotype in the *G. mellonella* model, since a reduced ability of the fungus to adhere to host cells can lead to an attenuated potential of tissue colonization. Our results can also be related to models where the EVs played protective roles, including those involving *Bacillus anthracis*, *Mycobacterium tuberculosis* or *Streptococcus pneumoniae.* In these organisms, previous treatment of mice with microbial EVs favored infection control ^69–71^. Our present data suggest that the protective roles of EVs can include mechanisms of attenuated virulence originated to an altered EV-regulated gene expression within infectious populations.

In conclusion, our results support recent findings showing that fungal EVs can be used as messaging compartments and directly influence growth, biofilm production and morphogenesis. The major players mediating yeast-to-hypha differentiation were characterized here, and further studies are necessary to determine whether these lipids in combination could be used to control candidiasis *in vivo*.

## Material and Methods

### Microorganisms and culture conditions

*C. albicans* ATCC strains 90028, 10231, SC5314 and the *Saccharomyces cerevisiae* strain RSY225 were cultivated in Sabouraud at 30 °C under agitation (150 rpm) for 48 h. *Histoplasma capsulatum* strain G-217B (ATCC 26032) was cultivated in HMM at 37 °C under agitation (180 rpm). Yeast cells of *C. albicans* were washed with phosphate-buffered saline (PBS) 3 times, enumerated and then transferred to medium 199 (M199) pH 4.0 or pH 7.0, RPMI-MOPS 0.165 M or RPMI supplemented with 10% fetal bovine serum (FBS) according to the experiments described below.

### Isolation and quantification of EVs

EVs were isolated following protocols developed by our group^30, 72^ with minor adjustments. Briefly, cultures (1l) from each fungal species were collected after 48 h of growth as described above. Cells and debris were removed by sequential centrifugation steps (4,000 and 15,000× *g*, 15 min, at 4 °C). An additional step of filtration using a 0.45 μm membrane filter (Merck Millipore) was used to remove any remaining yeast cells or debris. The supernatant was then concentrated using an Amicon ultrafiltration system (100-kDa cut-off, Millipore) to a final volume of approximately 20 ml. EVs were collected by centrifugation at 100 000× *g* for 1 h at 4 °C. Supernatant was discarded and the pellet enriched in EVs was washed twice with PBS pH 7.4 at 100 000× *g* for 1 h at 4° C. Alternatively, 300 μl containing 10^7^ yeasts of *C. albicans* were plate onto Petri dishes containing Sabouraud Dextrose Agar (SDA). After 24 h of growth cells were harvested using a cell scraper and transferred to 20 ml of PBS. Cells and debris were removed as described above, the supernatant was filtered using a 0.45 μm membrane filter and the EVs were collected by centrifugation at 100 000× *g* for 1 h at 4 °C. To isolate EVs from macrophages (RAW 264.7 macrophage-like; ATCC TIB-71) culture medium RPMI supplemented with 10% FBS (400 ml) with 70% cellular confluence was collected after 24 hours and centrifuged (300x *g* and 15,000× *g*, 15 min, at 4 °C). The same steps of filtration were used and the EVs were recovered after centrifugation at 100 000× *g* for 1 h at 4 °C. The presence of EVs was confirmed using Dynamic Light Scattering (DLS) analysis ^30^ and Nanotracking analysis ^46^. EVs were suspended in 200 μl of PBS and aliquots of 5-10 μl were used to perform the sterol quantification. Sterol content was measured using the quantitative fluorometric kit ‘Amplex Red Sterol Assay Kit’ (Molecular Probes, Life Technologies). Protein content was measured using the BCA kit (Pierce, Thermo Fisher Scientific, US). Additionally, aliquots of 5-10 μl were also plated onto Sabouraud-Agar plates to confirm that the preparation is sterile.

### Nanotracking analysis of *C. albicans* EVs

Nanotracking Analysis (NTA) of fungal EVs was developed on an LM10 nanoparticle analysis system, coupled with a 488-nm laser and equipped with an _S_CMOS camera and a syringe pump (Malvern Panalytical, Malvern, United Kingdom) following the conditions used by Reis and colleagues ^46^. All samples were 833-2,500-fold diluted in filtered PBS and measured within the optimal dilution range of 9 × 10^7^ to 2.9 × 10^9^ particles/ml. Samples were injected using a syringe pump speed of 50, and three videos of 60 s were captured per sample, with the camera level set to 15, gain set to 3, and viscosity set to that of water (0.954 to 0.955 cP). For data analysis, the gain was set to 10 to 15 and the detection threshold was set to 2 to 3 for all samples. Levels of blur and maximum jump distance were automatically set. The data were acquired and analyzed using the NTA 3.0 software (Malvern Panalytical).

### Fungal growth and biofilm Assays

Yeast cells of *C. albicans* (90028) were plated onto polystyrene 96-well plates (10^3^ yeast cells/well) containing Sabouraud or RPMI-MOPS (in accordance with the CLSI standard M27 guideline) in the presence or absence of distinct concentrations of EVs (0.5; 1 and 5 μg/ml, based on sterol content) and cultivated at 37 °C. Cellular density was monitored for up to 48 h by measuring the culture absorbance at 600 nm every 1 h using a Bioscreen C (Growth Curves USA). For Biofilm production and quantification, yeast cells were plated onto polystyrene 96-well plates containing M199 (10^5^ cells/well) in the presence of different concentrations of EVs (0.5; 1 and 5 μg/ml based on sterol content). PBS was used as a negative control. Biofilm was allowed to develop for 24 and 48 h at 37 °C and then quantified by Crystal Violet method. Briefly, after each time point the biofilm-coated wells were washed 3 times with PBS, air dried for 45 minutes and stained with 0.4% aqueous crystal violet (100 μl) for 10 minutes and washed 3 times with 200 μl of PBS. Then, 100 μl of absolute ethanol was added to the wells and the systems incubated for 5 minutes. The plate was then analyzed using a microplate reader (EL808, BioTek) at the wavelength of 540 nm ^73^. Tests were performed in triplicate. For Scanning Electron Microscopy yeast cells were washed three times in phosphate buffered saline (PBS) pH 7.2 ± 0.1 to remove planktonic cells and fixed in 2.5% glutaraldehyde type I, in 0.1 M sodium cacodylate buffer (pH 7.2 ± 0.1), for 1 h at room temperature. After fixation, cells were washed in 0.1 M sodium cacodylate buffer (pH 7.2 ± 0.1) containing 0.2 M of sucrose and 2 mM of MgCl_2_. Then, cells were dehydrated in ethanol (30, 50 and 70%, for 5 min, then 95% and 100% twice for 10 min), subjected to critical point drying in an EM CPD-300 (Leica, Wetzlar, Germany), covered with gold sputtering (10 ± 2 nm) and visualized in a Zeiss Auriga 40 (Zeiss, Oberkochen, Germany) microscope operated at 2 kV.

### Yeast-to-hyphae differentiation

Yeast cells of *C. albicans* (2.5×10^3^/well) were plated onto 96-well plates containing the hypha-inducing media M199 or RPMI supplemented with 10% Fetal Bovine Serum (FBS). To test whether the number of yeast cells would influence the EVs activity different suspensions of *C. albicans* were used (1.5×10^3^, 2×10^4^ and 3.5×10^4^ yeast cells/well). Fungal EVs were added at final concentration varying from 0.025 to 5 μg/ml (based on sterol content). For the lipid activity tests aliquots of 10 μl (equivalent to 5 μg/ml of EVs) were added to the wells. Plates were incubated at 37 °C and 5% CO_2_ for 4, 8 and 24 h. For each time point, the cells’ morphologies were visualized under an Observer Z1 microscope (Carl Zeiss International, Germany) and the images were collected and processed with ImageJ (NIH, US) and Photoshop (Adobe). Time-lapse video microscopies were performed after plating yeast cells of *C. albicans* strain 90028 (5 x 10^3^ yeast cells/well) onto 4-wells plate containing 1 ml of M199 medium. A final concentration of 5 μg/ml of *C. albicans* EVs was added to the wells. PBS was used as control. Samples were placed in a culture chamber with controlled temperature and CO_2_ environment (37 °C and 5%, respectively), attached to a Nikon Eclipse TE300 (Nikon, USA). Phase contrast images were captured every minute using a Hamamatsu C2400 CCD camera (Hamamatsu, Japan) connected with a SCION FG7 frame grabber (Scion Corporation, USA). Time-lapse images were taken every 1 min using a Nikon Eclipse TE300 microscope attached to a Digital Hamamatsu C11440-10C camera (Hamamatsu, Japan). Movies animations were generated using ImageJ 1.8 software (NIH, US).

### EV Lipid extraction

An aliquot containing 50 μg/ml of EVs were dried using a SpeedVac Vacuum Concentrator (Eppendorf), followed by lipid extraction with chloroform (C):methanol (M):water (W) (8:4:3; v/v/v) according to Nimrichter et al ^74^. The pellet containing non-extractable residues and precipitated protein was recovered by centrifugation. Partition was obtained after water addition to the system to reach the proportion C:M:W (8:4:5.6; v/v/v). The upper and lower phases and the protein content were dried separately using SpeedVac, suspended in DMSO and their activity tested as described below. The lower (LP) and upper phases (UP) contain lipids with higher and lower polarity, respectively. The interphase was also dried using SpeedVac and suspended in DMSO. Aliquots of each fraction (pellet, and lower and upper phases) equivalent to 5 μg/ml (based on sterol content) of EVs according to previous quantification were tested. The same protocol was used to separate the non-polar components of Sabouraud medium, and the lower phase was used as control.

### Thermostability of EV activity

To investigate the thermo stability of the *C. albicans* 90028 yeast EV activity the samples were heated for 15 min at 90 °C. Then a final concentration corresponding to 5 μg/ml (based on sterol content before heat treatment) of EVs was tested as described above.

### Long-term effect of EVs

Yeast cells of *C. albicans* (2.5×10^3^/well) were plated onto 96-well plates containing M199 and treated with 5 μg/ml (based on sterol content) of EVs. The *C. albicans* cell morphology was analyzed using an Observer Z1 microscope (Carl Zeiss International, Germany) after 4, 8 and 24 h. The yeast cells were collected after 24 h incubation in the presence of EVs and washed three times with PBS. Yeast cells were enumerated and a total of 2.5×10^3^/well was transferred to fresh M199, in the absence of EVs. These steps were repeated four times without addition of EVs to the cultures. Untreated yeast cells cultivated in M199 were used as control each 24 h. These steps were repeated four times.

### Hyphae-to-yeast differentiation

Yeast cells of *C. albicans* (2.5×10^3^/well) were plated onto 96-well plates containing M199 media and then incubated for 4 h to allow yeast-to-hypha differentiation. After this period 100% of the yeast cells differentiated to hyphae. Then, a final concentration of EVs (5 μg/ml, based on sterol content), EVs (5 μg/ml) and Tyrosol (200 μM) (Sigma-Aldrich, US) were added to the wells and the morphology accompanied for additional 4, 8 and 24 h as described above. PBS was used as control. Tyrosol stock was diluted in methanol.

### Yeasts Lipid Extraction

Yeast of *C. albicans, H. capsulatum* and *S. cerevisiae* (6×10^10^ cells) were suspended in a mixture of C:M (2:1, v/v), stirred for 16 h at room temperature and then clarified by centrifugation. Cells were reextracted with a mixture of C:M (1:2, v/v) under the same conditions. Lipid extracts were combined and dried using a rotavapor (Heidolph, Germany). The lipids were then suspended in a mixture of C:M:W (8:4:5.6; v/v) and vigorously mixed. The lower phase was dried using a rotavapor and suspended in 2.5 ml of chloroform. Aliquots were dried using Speedvac (Eppendorf, Germany) and suspended in DMSO. A yeast-to-hypha differentiation assay was performed to determine the minimal amount of the lower phase from *C. albicans* (20; 4; 2; 1; 0.5; 0.25; 0.125 and 0.05 mg/ml) was able to block the differentiation. Aliquots of 1 mg/ml were used to the transcriptomic assay.

### Size-Exclusion Chromatography (SEC)

To exclude the presence of free-contaminants in EV preparations we used size-exclusion chromatography (SEC) according to Menezes-Neto and colleagues ^75^ with minor adjustments. Briefly, Sepharose 4B (final volume of 10 mL) was packed in a 10 mL syringe and then equilibrated with PBS-citrate 0.36% (w/v). The EVs obtained after ultracentrifugation were suspended in a final volume of 500 μL and applied to the column. A total of 30 fractions of 500 μL were collected immediately after column loading using PBS-citrate as eluant. EVs aliquots of 5-10 μl were used to perform the sterol and protein quantification as described previously. A yeast-to-hypha differentiation assay was performed to determine the activity of each fraction. To avoid additional steps to concentrate the samples the lipids from each fraction were extracted as described above and the LP normalized according to the sterol content. A final concentration of 5 μg/mL of sterol was used. For the fractions where sterol was not detected 400 μL of each fraction was used for lipid extraction and solvent partition. The LP was dried down in a speed vac and suspended in 2 μL of DMSO. An aliquot of 1 μL was used to test the inhibitory activity.

### Reverse-phase thin-layer chromatography (RP-TLC)

EVs or LP lipids were suspended in C:M (1:1, v/v) and their concentration normalized by sterol content or the total weight, respectively. Aliquots of 10 μL were spotted into reverse-phase TLC (RP-TLC) plates (silica gel 60, RP-18, F_254_s, 1 mm, Merck, Germany). The plates were developed in chambers pre-saturated for 10 min using acetone as a solvent system. After development, plates were dried at room temperature and the bands were visualized after spraying with a solution of 50 mg ferric chloride (FeCl_3_) in a mixture of 90 ml water, 5 ml acetic acid, and 5 ml sulfuric acid^76^. After spraying, plates were heated at 100 °C for 3 to 5 min. Farnesol was used as standard. Under these conditions, farnesol appears as a purple band. Alternatively, the developed plates were incubated in a saturated iodine chamber where the lipids appear as yellow bands. The bands were scraped from the TLC and placed into microcentrifuge tubes. Lipids were suspended in C:M (1:1, v/v), filtered using a PVDF membrane with 0.22 μm to a new microcentrifuge tube. Samples were dried down using a speed vac, suspended in DMSO and normalized by lipid weight. For the lipid activity tests aliquots of 10 μl (equivalent to 5 μg/ml of EVs) were added to the wells containing yeasts of *C. albicans* in M199 medium. Plates were incubated at 37 °C and 5% CO_2_ for 4, 8 and 24 h. For each time point, the cells’ morphologies were visualized under an Observer Z1 microscope (Carl Zeiss International, Germany) and the images were collected and processed with ImageJ (NIH, US) and Photoshop (Adobe).

### Agar invasion assay

Hyphal growth was stimulated using M199 medium. Yeast cells of *C. albicans* (90028) were treated or not with EVs (5 μg/ml, based on sterol content) overnight and a suspension of yeast cells 10 μl containing 10^5^ cells was plated onto M199 Agar plates. Plates were then incubated at 37 °C and growth was monitored daily for hypha formation. After 10 days of incubation colonies were washed in a stream of water using the same flow rate, water temperature and period of time as described by Cullen^56^. Colony images were taken with an Optiphase microscope coupled with a digital camera (AmScope, US) and processed using Adobe Photoshop.

### Farnesol extraction

The *C. albicans* yeast 90028 EV sample (19.1 μg total sterol) was resuspended in 400 µl methanol (HPLC grade, Fisher Scientific), vortexed for 2 min and centrifuged for 15 min, at 14,000 x g, at room temperature. The supernatants were transferred to a 2-mL PTFE-lined cap glass tube (Supelco, Cat#27134, Sigma-Aldrich). The ensuing pellet was redisolved in 400 µl methanol and the previous steps were repeated. The two resulting supernatants were pooled and dried under nitrogen gas in Supelco PTFE-lined cap glass vials. For farnesol extraction, the yeast 90028 EV sample was partitioned by adding 500 µL HPLC-grade water (Fisher Scientific) and 500 µL ethyl acetate (Sigma-Aldrich, cat # 270989) and vortexed for 2 min. After phase separation, the upper (organic) phase was collected, transferred to a fresh PTFE-lined cap glass vial, and dried under nitrogen gas. Samples were then redisolved in 20 µL methylene chloride (Optima grade, Fisher Scientific), transferred to Verex PTFE-lined/silicone cap vials (Phenomenex, µVial, i3 Qsert), just prior to gas chromatography-mass spectrometry (GC-MS) analysis.

### Farnesol analysis by GC-MS

A standard of farnesol isomer mixture (catalog # 93547, Supelco, Sigma-Aldrich) was diluted in methylene chloride (Optima grade, Fisher Scientific) to a concentration of 22.5 picomole/µL to determine the transition ions to be used in the single-reaction monitoring (SRM) method. Chromatographic separation was performed by injecting 1 µl of farnesol standard onto a Thermo TSQ™ 9000 Triple Quadrupole GC-MS/MS System (Thermo Scientific) set to splitless, MS transfer line at 250°C, with an AI/AS1310 series autosampler (Thermo Scientific), equipped with a Zebron (ZB-FFAP, 30 m x 0.32 mm x 0.25 µm) column, previously indicated for separation of farnesol isomers ^52^. Initially, a full scan was run to determine retention time (RT) and underivatized farnesol limits of detection, prior to farnesol SRM confirmation and full scan analysis.

The lowest detectable level of underivatized farnesol isomers from the standard, using the automated SRM (AutoSRM) mode on the TSQ 9000 Triple Quadrupole MS (Thermo Scientific) with an electron ionization (EI) source temperature at 200°C, was 0.0045 picomole/μL. However, subsequent analyses were performed with a concentration of farnesol standard of 4.5 picomole/μL, in which the detection of all four isomers (Z,Z, Z,E, E,Z, and E,E) of farnesol was reproducible, and the signal of the least abundant isomer (Z,Z) was at least ∼10% of the most abundant isomer (E,E) and ∼6-fold higher than the background. Using the AutoSRM mode on the TSQ 9000 dashboard feature, the ion fragmentation was optimized at their respective RT with the top 3 transition ions (81.1 to 79.1 at 8 collision energy (CE), 69.1 to 41 at 6 CE, and 41.1 to 39.1 at 8 CE) were chosen and imported into Chromeleon v7.2.9. Images of the SRM chromatograms of the farnesol standard and the sample extracted from *C. albicans* yeast 90028 EVs were transferred from Chromeleon and PowerPoint for figure generation.

### Quantitative farnesol and farnesol-like molecules by GC-MS

Farnesol quantitation was measured by building a standard curve with d6-E,E-farnesol (catalog # SKU700294O, Avanti Polar Lipids, Alabaster, AL) at concentrations of 0.1 ng/µL, 1 ng/µL, 2.5 ng/µL, and 5 ng/µL. Peak areas of the standard and samples were imported into GraphPad Prism 9.0.0 for formula and R^2^ generation, and graph.

### RNA isolation, purification, and sequencing

The cells were treated with EVs (5 μg/ml, based on sterol content), Farnesol (FOH) (150 μM) and lipidic lower phase (LP) (1 μl) for 18 h and the RNA was extracted by using the mirCury cell and plant isolation kit (QIAGEN). A final number of 10^8^ treated cells were lysed using 500 μL of glass beads and the 600 μL of the Lysis buffer provided with the kit. Three biological replicates were obtained for each condition. The concentration and RNA integrity were analyzed using Qubit (Life Technologies) and bioanalyzer 2100 (Agilent Technologies, CA). The RNA-seq libraries were constructed using the TruSeq Stranded mRNA kit (Illumina). According to the manufacture instructions. RNA sequencing was performed using an Illumina HiSeq 2500 equipment at the Life Sciences Core Facility (LaCTAD) from State University of Campinas (UNICAMP, Brazil).

### Transcriptomic analysis

The data analysis was performed using CLC Genomics Workbench 10.0.1 (Qiagen). Quality trimming and adapter trimming were performed using default parameters. Reads were mapped to the *C. albicans* strain SC5314 reference genome (ASM18296v3). Differential gene expression profiling was carried out using the negative binomial distribution. The false-discovery rate adjusted p-value (FDR) procedure was used for multiple-hypothesis testing correction. Genes with FDR-adjusted *P* value (less than or equal to 0.05) and expression fold change of more than 3 or less than 3 were considered to be differentially expressed. Functional enrichment analysis was performed on the differentially expressed genes to identify overrepresented Gene Ontology (GO) biological processes using the CLC Genomics Workbench and the David annotation tool ^77^. The enriched metabolic pathways were analyzed at the Kyoto Encyclopedia of Genes and Genomes (KEGG) ^78^ and using David annotation tool ^77^. To analyze the interactions between the transcripts we used the Protein-protein tool association STRING (Search Tool for the Retrieval of Interacting Genes) database ^79^. The parameters we used were: High confidence – minimum required of 0.700, interaction sources based on experiments, network edge based on confidence.

Data availability. The RNA sequencing data were deposited into the Sequence Read Archive (SRA) database from NCBI under accession number PRJNA203278.

### Galleria mellonella infection

Larvae of wax moth *G. mellonella* was used to investigate virulence of EV-treated yeast cells ^80^. *C. albicans* yeast cells (2.5×10^3^) were treated with EVs (5 μg/ml, based on sterol content) in M199 for 24 h. Yeast cells were then washed in PBS and enumerated. As a control yeast cells cultivated in Sabouraud were used. A final volume of 10 μl containing 2×10^5^ yeast cells was injected into the last right proleg of the larvae (200-300 mg each) using a Hamilton syringe. An additional control included larvae injected with 10 μl PBS alone. The injected larvae were kept at 37 °C and the number of deceased subjects per group was monitored daily.

### Statistical analysis

All statistical analyses were performed using GraphPad Prism 6, version 5.02 for Windows (GraphPad Software).

## Acknowledgements

This work was supported by grants from the Brazilian agency Conselho Nacional de Desenvolvimento Científico e Tecnológico (CNPq, grants 405520/2018-2, 440015/2018-9, and 301304/2017-3 to M.L.R.; 311179/2017-7 and 408711/2017-7 to L.N.; 311470/2018-1 to AJG), FAPERJ (E-26/202.809/2018 to LN, E-26/202.696/2018 and E-26/202.760/2015 to AJG), FIOCRUZ (grants VPPCB-007-FIO-18 and VPPIS-001-FIO18) and Coordenação de Aperfeiçoamento de Pessoal de Nível Superior (CAPES, Finance Code 001). J.D.N. was supported in part by NIH R21 AI124797. We thank the staff of the Life Sciences Core Facility (LaCTAD) from State University of Campinas (UNICAMP) for the RNA-seq facility. This work was also supported in part by NIH grants AI136934, AI116420, and AI125770, and in part by Merit Review Grant I01BX002924 from the Veterans Affairs Program to MDP. ICA was partially supported by grant # 5U54MD007592 from the National Institute on Minority Health and Health Disparities (NIMHD), a component of the National Institutes of Health (NIH). We thank the Biomolecule Analysis and Omics Unit at the Border Biomedical Research Center, UTEP, supported by NIMHD grant # 5U54MD007592 (to Robert A. Kirken), for the access to the GC-MS and other instruments. MTM received a Ph.D. fellowship from the Science Without Borders Program, sponsored by the Capes Foundation within the Ministry of Education, Brazil (grant n. BEX 013622/2013-07).

## Competing Interests Statement

Dr. Maurizio Del Poeta, M.D. is a Co-Founder and Chief Scientific Officer (CSO) of MicroRid Technologies Inc.

