## Supplemental Figures for "Extracellular vesicles regulate yeast growth, biofilm formation, and yeast-to-hypha differentiation in *Candida albicans*"

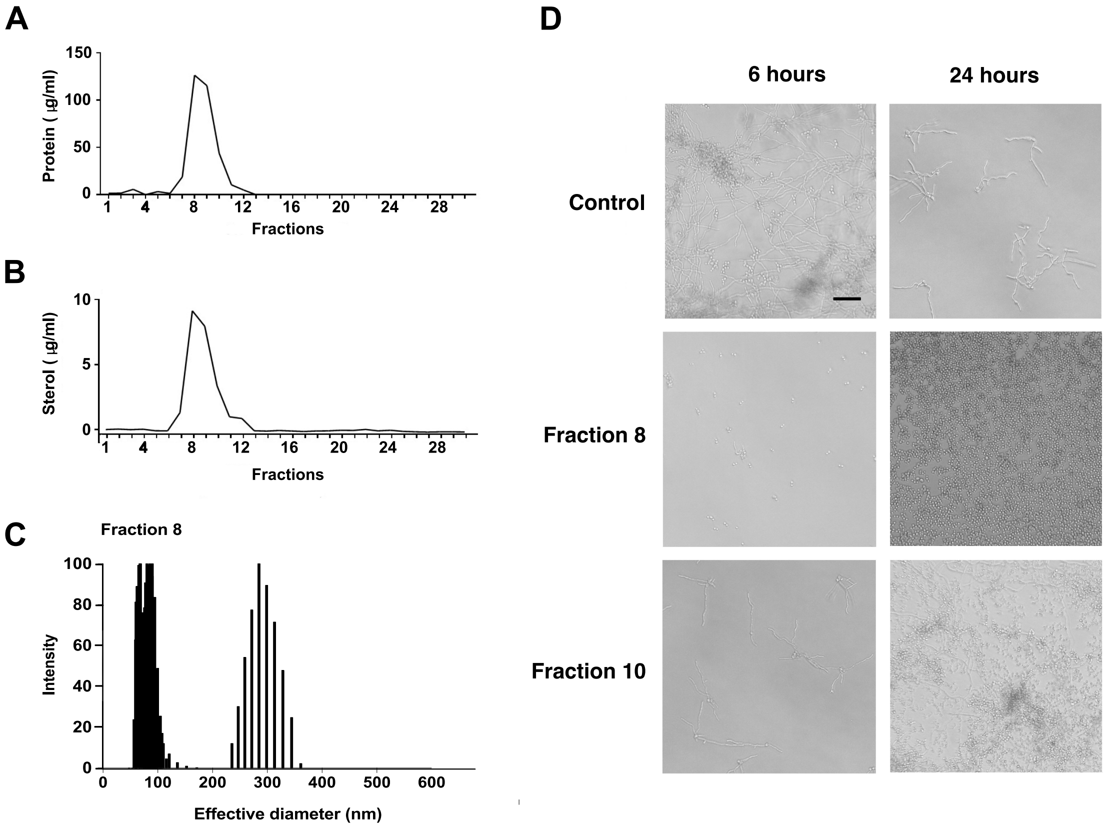


**Figure S1. Size-exclusion chromatography (SEC) and DLS of EVs from *C. albicans.*** The *Ca* EVs released into medium was enriched by SEC, using Sepharose CL-4B, and the 500-µL fractions collected as described in Methods. Each fraction was quantified according to its protein and sterol content. Fraction 8 displayed two populations of EVs with effective diameters usually found in *Ca* EVs. Yeasts of *C. albicans* (90028) were inoculated into M199 in the presence of each fraction to evaluate their yeast-to-hypha inhibitory activity. Fractions containing EVs were normalized according to the sterol content and 5 µg/ml were used. Fractions with no protein or sterol were also tested and 80% of the total content was used. None of these fractions showed any inhibitory activity. Effect was visualized after 6 and 24 hours. PBS was added alone as a control. Bar, 30 µm.

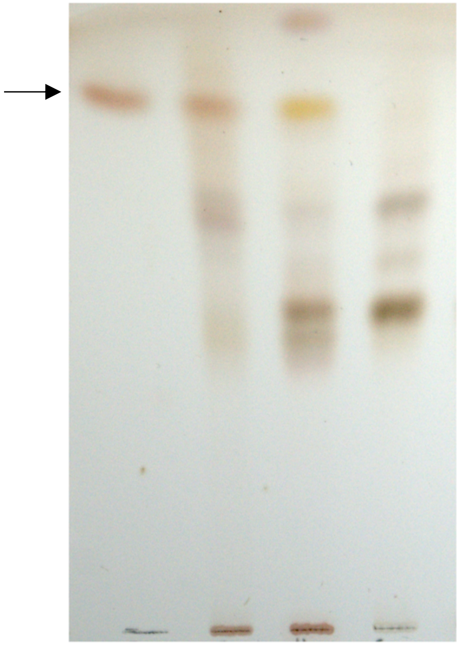


**Figure S2. Major lipids from *C. albicans, H. capsulatum* and *S. cerevisiae* resolved by TLC.** LP enriched in non-polar lipids from each fungal species was obtained as described in Methods. *C. albicans* (C*a*), *H. capsulatum* (*Hc*), and *S. cerevisiae* (*Sc*) lipid samples were resolved by reverse-phase TLC using acetone as solvent system and stained with ferric chloride-sulfuric acid reagent. Farnesol (FOH), indicated by an arrow, was used as standard.

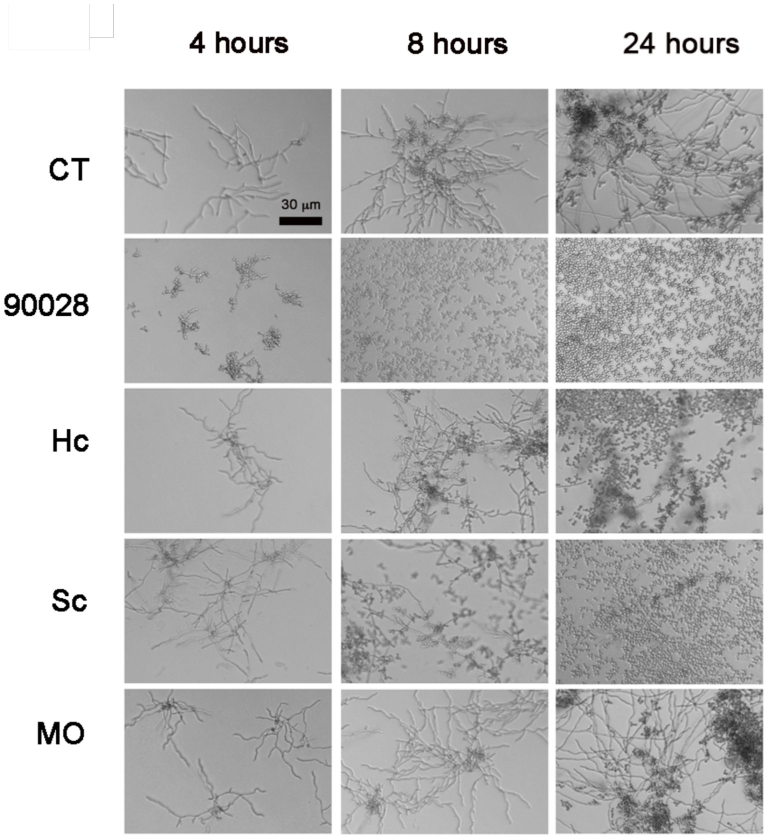


**Figure S3. Effect of EVs from different fungal species and macrophages over yeast-to-hypha differentiation.** Yeasts of *C. albicans* (90028) were inoculated into M199 in the presence or absence of EVs from *H. capsulatum* (*Hc*), *S. cerevisiae* (*Sc*), and macrophage-like RAW cell line (MO) at the equivalent concentration of 5 µg sterol/ml. Effect was visualized after 4, 8, and 24 h. PBS was added alone as a control (CT). Bar, 30 µm.

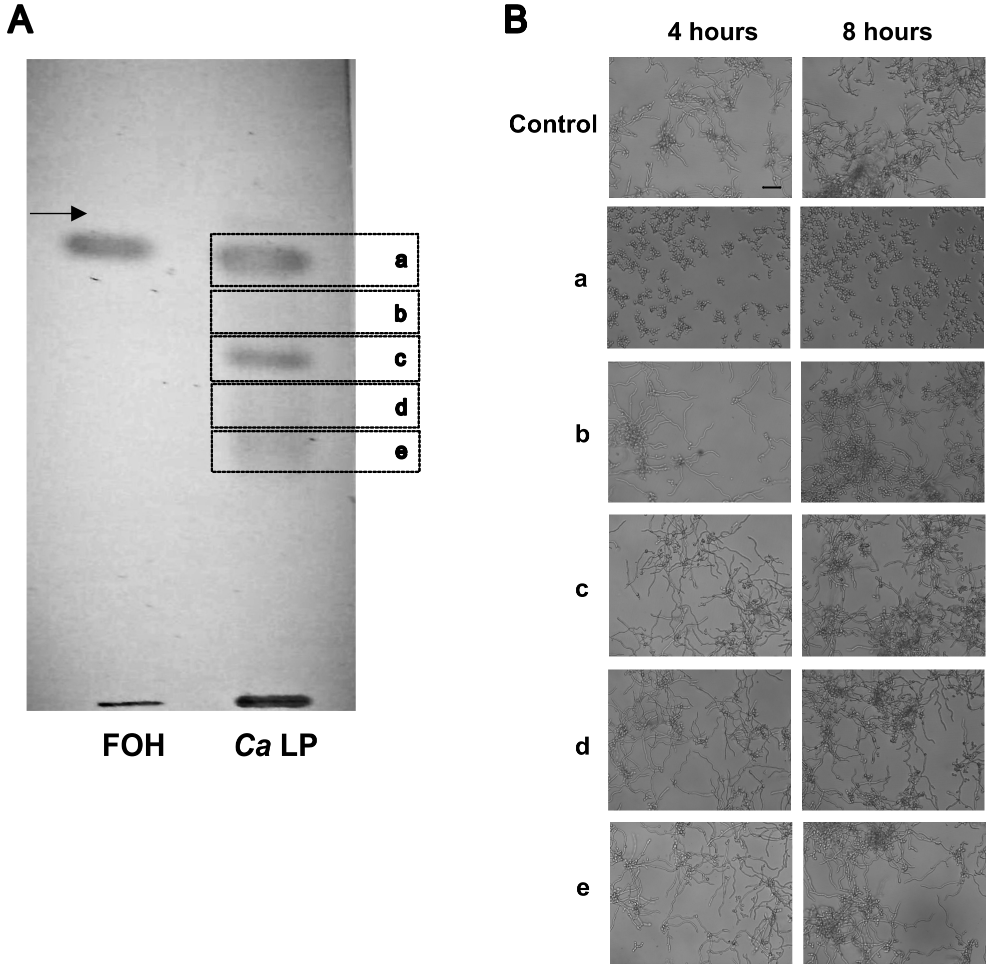


**Figure S4. Effects of bands isolated by preparative reverse-phase thin layer chromatography (RP-TLC).** (A) Lipids from LP of *C. albicans* strain 90028 (*Ca* LP) were resolved by RP-TLC and the bands (a-e) stained with $I_2$ vapor were scraped and eluted from the silica (see Methods). Farnesol (FOH), indicated by an arrow, was used as standard. (B) Yeasts of *C. albicans* (90028) were inoculated into M199 in the presence or absence of each eluted RP-TLC band (suspended in DMSO as described in Methods) Effect was visualized after 6 and 24 h. PBS-DMSO was added alone as a control. Bar, 30 μm.

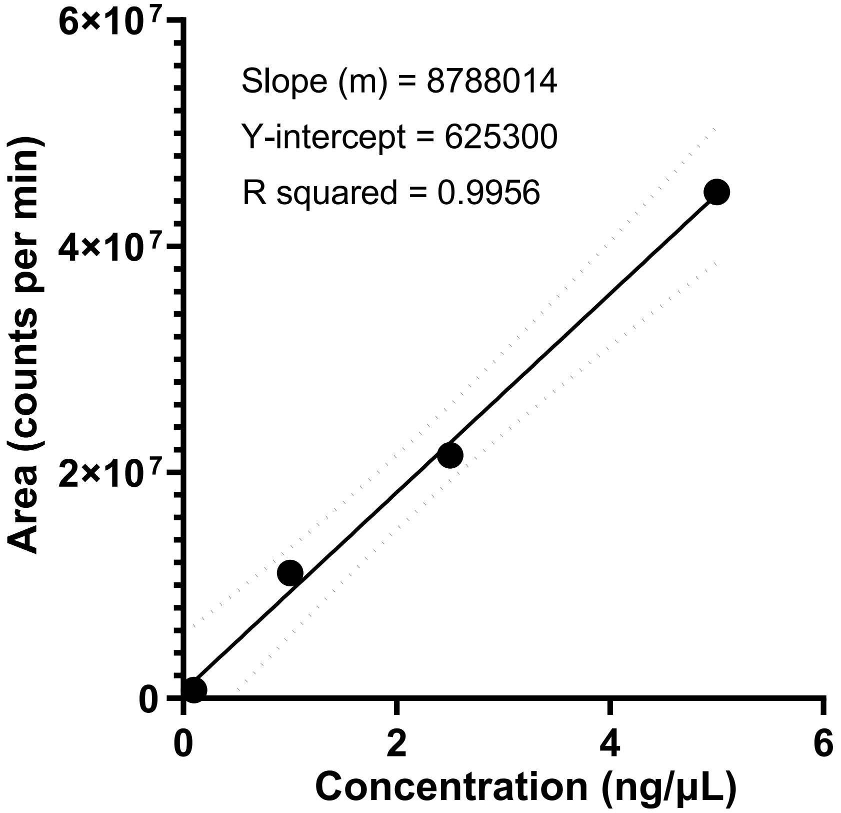


**Figure S5. Standard calibration curve of d6-farnesol E,E isomer.** d6-Farnesol E,E isomer (Avanti Polar) was diluted to concentrations of 0.1, 1.0, 2.5, and 5.0 ng/µL. Peak areas were imported to GraphPad v9.0.0 for graph and linear regression generation. The 95% confidence interval (discontinued line) of the curve is indicated.

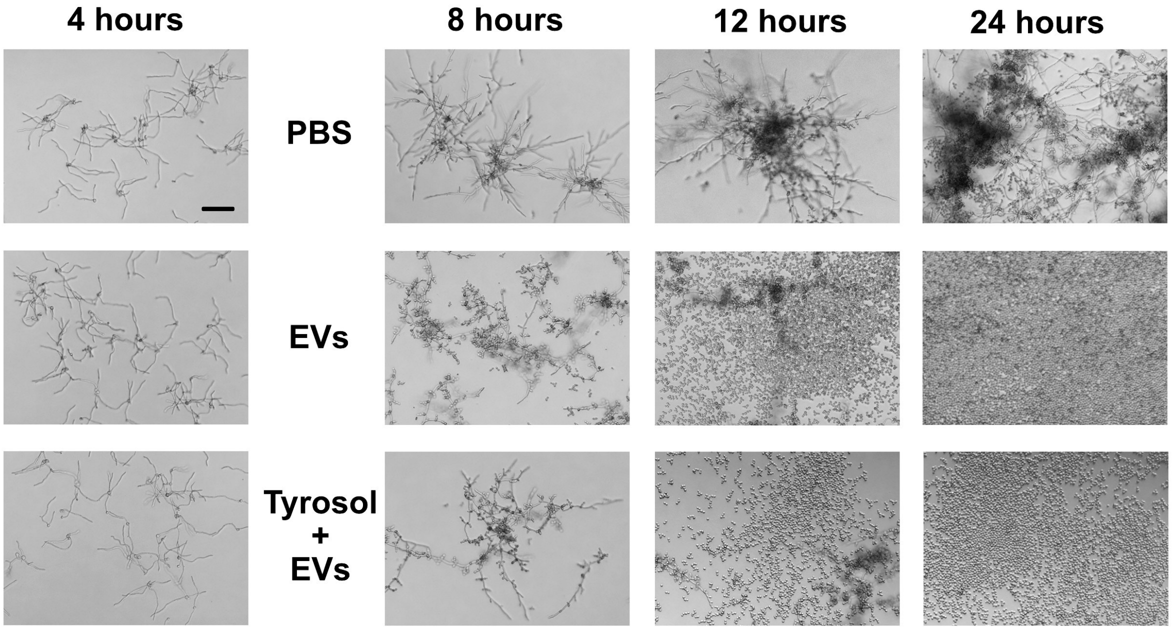


**Figure S6 –*C. albicans* EVs reversed the yeast-to-hypha differentiation.** Yeasts of *C. albicans* (90028) were inoculated into M199 and incubated for 4 hours to induce hyphae formation. Then PBS (control), *Ca* EVs (5 µg/ml) or a combination of EVs (5 µg/ml) and tyrosol (200 µM) were added to the wells. Cell morphology was visualized after 8, 12 and 24 hours. Bar 30 µm.

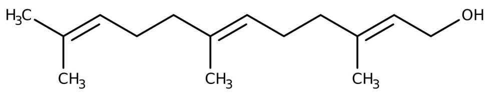


**Farnesol**

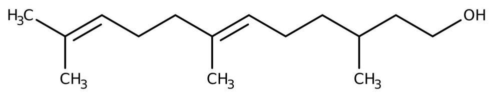


**6,10-Dodecadien-1-ol**

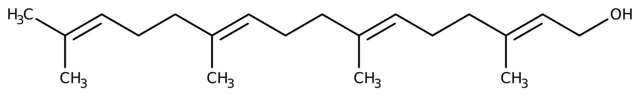


**Geranylgeraniol**

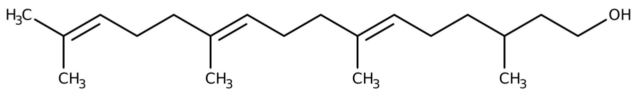


**6,10,14-Hexadecatrien-1-ol**

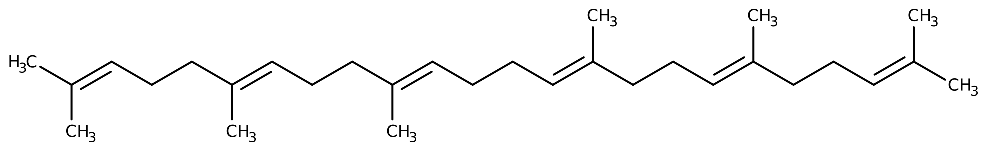


**Squalene**

**Figure S7. Major terpenoids or isoprenoids identified in *C. albicans* EVs and lipid extracts.**
